# A Multiscale approach reveals the molecular architecture of the autoinhibited kinesin KIF5A

**DOI:** 10.1101/2024.01.18.576271

**Authors:** Glenn Carrington, Uzrama Fatima, Ines Caramujo, Tarek Lewis, David Casas-Mao, Michelle Peckham

## Abstract

Kinesin-1 is a microtubule motor that transports cellular cargo along microtubules. KIF5A is one of three kinesin-1 isoforms in humans, all of which are autoinhibited by an interaction between the motor and an IAK motif in the proximal region of the C-terminal tail. The C-terminal tail of KIF5A is ∼80 residues longer than the other two kinesin-1 isoforms (KIF5B and KIF5C) and it is unclear if it contributes to autoinhibition. Mutations in KIF5A cause neuronal diseases and could affect autoinhibition, as reported for a mutation that skips exon 27, altering its C-terminal sequence. Here, we combined negative-stain electron microscopy, crosslinking mass spectrometry (XL-MS) and AlphaFold2 structure prediction to determine the molecular architecture of the full-length autoinhibited KIF5A homodimer, in the absence of light chains. We show that KIF5A forms a compact, bent conformation, through a bend between coiled coils 2 and 3, around P687. XL-MS of WT KIF5A revealed extensive interactions between residues in the motor, between coiled-coil 1 and the motor, between coiled-coils 1 and 2, with coiled coils 3 and 4, and the proximal region of the C-terminal tail and the motor in the autoinhibited state, but not between the distal C-terminal region and the rest of the molecule. While negative stain electron microscopy of exon-27 KIF5A splice mutant showed the presence of autoinhibited molecules, XL-MS analysis suggested that its autoinhibited state is more labile. Our model offers a conceptual framework for understanding how mutations within the motor and stalk domain may affect motor activity.

## Introduction

Kinesins are ATP-driven motor proteins that move along microtubules in eukaryotic cells. They are encoded by the 45 kinesin (KIF) genes in humans and can be classified into 14 separate classes based on sequence similarity (1). The majority of these classes comprise a kinesin with an N-terminal motor domain, which contains the nucleotide and microtubule binding sites, followed by a coiled-coil tail, which dimerises the kinesin and C-terminal cargo binding domains (N-kinesins) (Fig. 1A). The cargo binding domains determine to which cargo the kinesins bind and traffic (2, 3). The N-kinesins move towards the plus (fast growing end) of microtubules, trafficking cargo away from the cell body towards the cell periphery (anterograde transport).

**Fig. 1.**
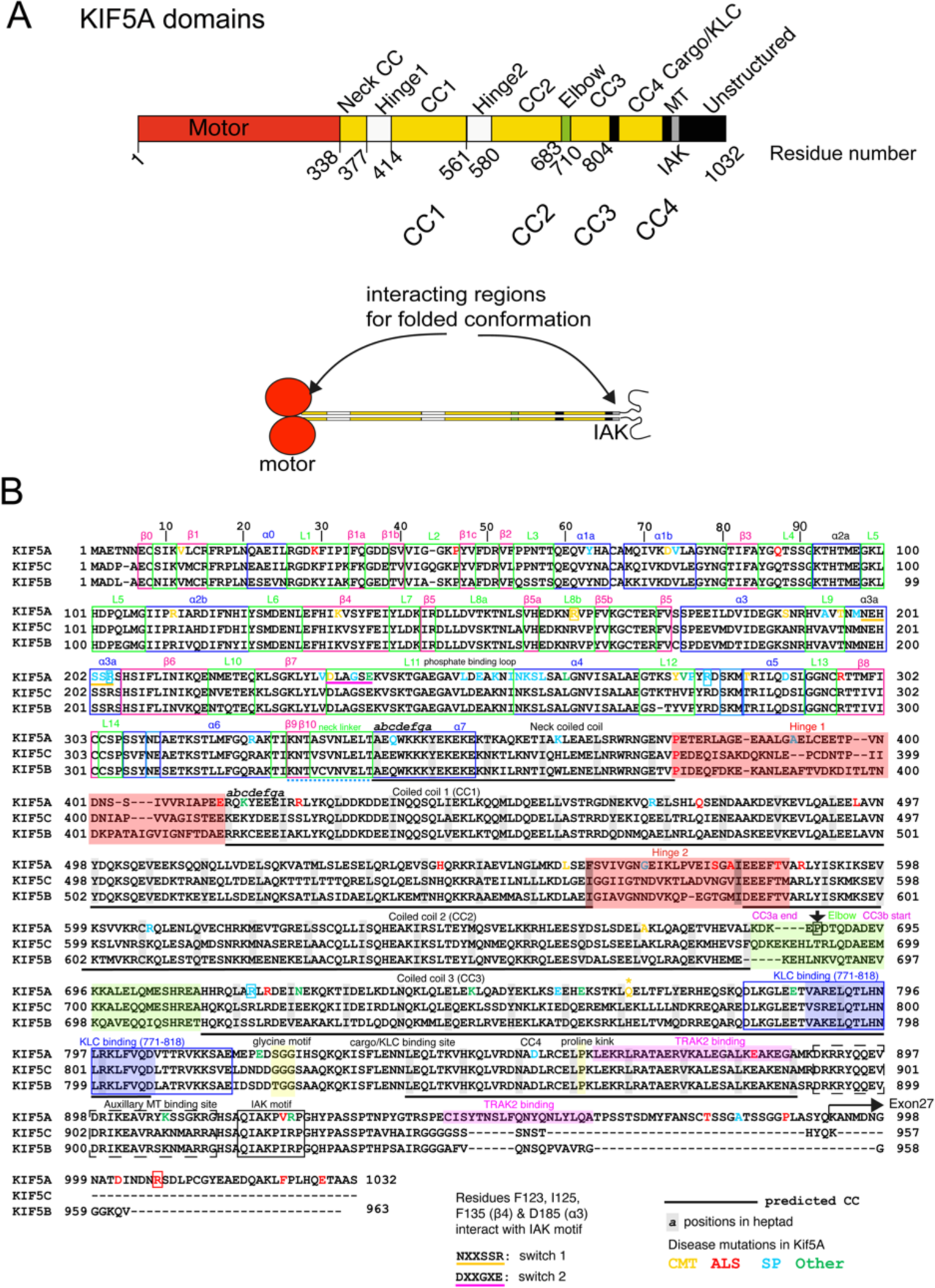
Overall organisation of human KIF5A. **A:** KIF5A is comprised of various domains; motor, coiled coils and C-terminal unstructured region. The IAK motif in the C-terminal region of the tail interacts with the motor domain in the shutdown molecules as indicated (18). **B:** Sequence alignment of human KIF5A (Q12840), KIF5B (P33176) and KIF5C (O60282), showing the positions of secondary structure within motor domain, predicted coiled-coil and C-terminal tail domains. The positions of mutations for Spastic Paraplegia type 10 (SP), Amyotrophic lateral sclerosis (ALS) and Charcot Marie Tooth Disease (CMT) that affect KIF5A are shown. Mutations that affect splicing of exon 27 result in altered sequence from residue 992 onwards as indicated by the arrow. Mutations were obtained from the human genome database (HGMD). The position of a hinge in KIF5B reported to be the position of folding of the molecule (between ‘CC3a and CC3b’) is also shown in magenta (19). For clarity, the neck coil is also termed CC1 (19) or C0 (18) in other previous studies.

Kinesin-1 (conventional kinesin, or KIF5) is an N-kinesin comprised of a homodimer of two heavy chains (4). Its C-terminal tail also binds two light chains to form heterotetramers, and the light chains act as cargo adaptors (5). Three genes (KIF5A, KIF5B and KIF5C) encode the heavy chains and four genes (KLC1-4) encode the light chains. The heavy chains can also bind to and direct cargo transport directly in a light chain independent manner (5), for example, directly binding to and sliding microtubules through its tail (6), ooplasmic streaming in *Drosophila* (7), binding to TRAK1 or 2 to traffic mitochondria (8, 9) and ER movement (10). All three isoforms are primarily involved in trafficking organelles, proteins and RNA (5). KIF5B is ubiquitously expressed while KIF5A and C are predominantly expressed in neurones (11).

All three kinesin-1 isoforms form an autoinhibited state, in which the kinesin is unable to bind to cargo and thus kinesin-1 based transport is regulated. It was originally shown that a conserved isoleucine-alanine-lysine (IAK) motif in the C-terminal region of the heavy chain interacts with and inhibits motor domain kinesin-1 ATPase activity and reduces its affinity for microtubules (12, 13) (Fig. 1A). Structural studies using Cryo-EM and crosslinking mass spectrometry of motor domains crosslinked to tail peptides demonstrated that the IAK motif interacts directly with switch-1 in the motor to inhibit its activity (14). However, only one of the two IAK motifs interacts with both motor domains in this inhibited state (15, 16). Moreover, mutations in the IAK motif can increase the association of KIF5B with microtubules but do not increase processive movement along them (17). Thus, the IAK motif by itself may not be sufficient to regulate kinesin activity.

Until recently, the position in the sequence at which kinesin bends to enable the interaction of the tail domains with the motor domain in the autoinhibited molecule was unclear. Recent negative stain electron microscopy (nsEM) studies of KIF5C in complex with KLC1 combined with Alphafold modelling demonstrated that this bending likely occurs in a region termed the elbow (at around residue 690) (18). The elbow lies between coiled-coil 2 and coiled-coil 3, in a region that is 19 residues long starting at residue 683 (human KIF5C) (Fig. 1A,B). A further study of KIF5B and KIF5C, which combined nsEM with crosslinking followed by mass spectrometry, reported that kinesin bends at a break in coiled-coil 3 (CC3, as defined in (19)) between CC3a and CC3b. This break point is located at around residue 690, and thus is in approximately the same region as the elbow (19). Both KIF5C and KIF5B homodimers formed autoinhibited molecules. The addition of light chains to form heterotetramers did not affect the folding pattern, but the presence of light chains likely stabilised the autoinhibited state (19). The position of this bend has not yet been demonstrated for KIF5A.

While the amino acid sequences of KIF5A, KIF5B and KIF5C are highly conserved (∼70% identity), the sequences of the C-terminal regions distal to the IAK motif are highly variable (Fig. 1B). The C-terminal tail of KIF5A is about 70 residues longer than that of KIF5B and KIF5C, is likely to be intrinsically disordered (20) and its function is unclear. KIF5A has a unique role in trafficking mitochondria (21). To achieve this, KIF5A is directly coupled to TRAK2 (Trafficking Kinesin Protein 2) via sites mapped to the C-terminal tail (22) and TRAK2 activates KIF5A independently of light chains (8). TRAK2 activation of KIF5A may be distinct from the synergistic regulation of kinesin-1 activity via associated light chains in which the light chains stabilise the autoinhibited state formed by the heavy chain on folding up and are important in helping to activate kinesin through binding to cargo (17–19).

Mutations that result in the aberrant splicing of exon 27 cause amyotrophic lateral sclerosis (ALS). These mutations replace the sequence of the last 34 residues of KIF5A with a novel 39 amino acid sequence, which has been suggested to disrupt its autoinhibition (23). It is not entirely clear why this is the case, but it has been suggested that the altered sequence changes the overall charge of this region from positive to negative, which may in turn affect its interaction with the downstream negatively charged region containing the IAK motif and interfere with its ability to interact with the motor domains in the autoinhibited state (23). It has also been suggested that this mutation is a toxic gain of function mutation that aggregates rather than activates KIF5A and interferes with trafficking of mitochondria (24–26).

Understanding how KIF5A is autoinhibited is not only important for understanding its role in trafficking but also in disease. Disease causing mutations in KIF5A are far more common than those in KIF5B and KIF5C (Fig. 1B). They are responsible for a range of neuropathies including Spastic Paraplegia type 10 (SP), Amyotrophic lateral sclerosis (ALS) and Charcot Marie Tooth Disease (CMT) (27, 28) (Fig. 1B, Supplemental Table 1).

Here, we set out to use nsEM in combination with crosslinking mass spectrometry (XL-MS) of purified full-length KIF5A in its autoinhibited state to determine if it folds up in a similar way to that reported for KIF5B and C, and what, if any, role the extended C-terminal tail might have in autoinhibition. We performed these experiments for the homodimeric KIF5A in the absence of light chains. We used Alphafold2 to build a model of the autoinhibited molecule, using the nsEM and XL-MS data to help verify the model. We also tested the effect of the exon 27 splice mutation on formation of the autoinhibited molecule using nsEM and determined if there were any changes to crosslinking of KIF5A in the autoinhibited molecule using XL-MS. Finally, we tested the effects of the exon 27 splice mutation and 3 further missense mutations in the C-terminal tail on the location of eGFP-KIF5A in cells.

## Results

### Full-Length KIF5A expressed and purified from Sf9 cells forms an autoinhibited dimer

Full-length human KIF5A heavy chain was expressed using the baculovirus/Sf9 system in the absence of KLC. The molecular mass of the purified protein had the expected molecular weight of approximately 120 kDa (Fig. 2A). No endogenous light chains (molecular mass of KLC1-4 range from 55 to 70kDa) or adaptor proteins such as TRAK2 (101kDa) from the Sf9 cells appeared to copurify with the heavy chain. Mass photometry of molecules in low salt buffer (150mM KCl), in conditions where KIF5A is expected to be autoinhibited, revealed that the majority of the purified protein had a molecular weight of 227 kDa (Fig. 2B) consistent with dimer formation by KIF5A. A second smaller peak was also observed at twice this molecular weight (474 kDa), which is likely to represent tetramers of KIF5A (Supplemental Fig. 1A,B). Tetramer formation in the absence of light chains has been reported previously for this kinesin isoform (17).

**Fig. 2:**
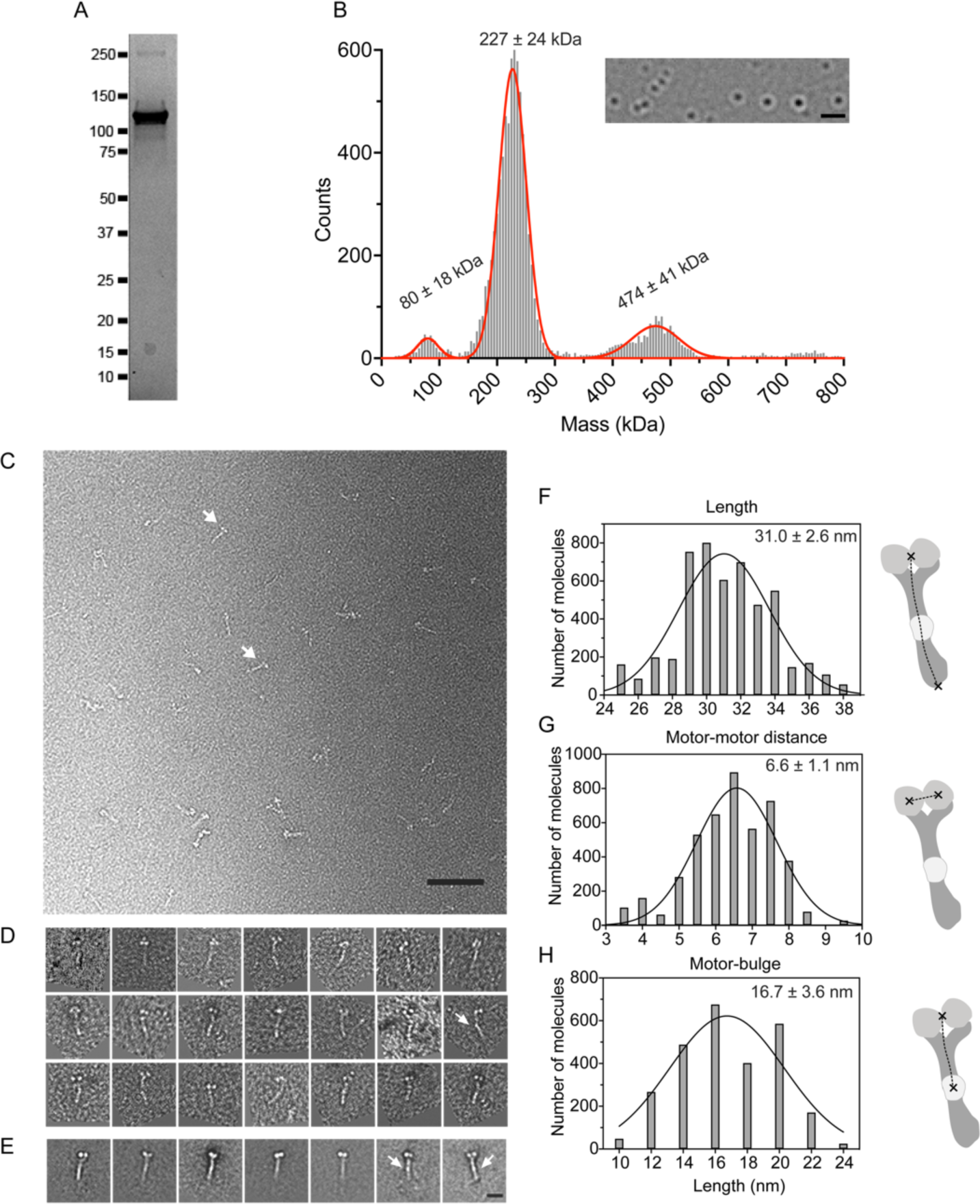
Purification, mass photometry and negative stain EM analysis of purified KIF5A. **A:** SDS gel of the purified KIF5A from Sf9 cells. The molecular mass of the purified protein is ∼120kDa. **B:** Mass Photometry results for KIF5A. Numbers above each peak show the mean molecular mass for 3 measurements ± S.D. The image insert shows the type of image obtained in this experiment. using the rolling background subtraction method as described (50). Scale bar 1 µm. **C:** A representative field of view of autoinhibited KIF5A molecules imaged by nsEM. Arrows indicate individual molecules. Scale bar 200nm. **D:** examples of individual KIF5A molecules imaged by nsEM. **E:** image classes of KIF5A molecules. Arrows indicate a ‘bulge’ midway along the coiled-coil region in D and E. Scale bar is 10nm. Measurements of the lengths of the stalk, distance between the motor domains and between the motor domains and the bulge halfway along the molecule, are shown in **F,G and H** respectively. Mean ± S.D for these measurements are as shown.

KIF5A was imaged by nsEM under conditions in which the molecule forms the autoinhibited state (2 mM Mg.ATP and low salt (150 mM KCl)) (Fig. 2C). The raw particles were characterised by the presence of 2 globular densities, which we attribute to the motor domains based on their size, shape and morphology. A long thin structure, likely to be formed by the coiled-coil tail, extends from the two globular motor domains, and an additional small additional density appears halfway along this structure in some views (Fig. 2D). The raw images showed that the majority of the molecules were in the autoinhibited state. A small number of extended molecules, or molecules that might be tetrameric were also occasionally observed (Supplemental Fig. 1). Individual dimeric molecules were picked, aligned and used to generate class averages (Fig. 2E), which showed two globular motor domains, followed by a long region of tail (the ‘stalk’). The additional density half-way along the stalk was easily visible in some classes (Fig. 2E).

The stalk length, the distance between the motor domains and the end of the stalk, was 31 ± 2.6 nm (mean ± SD) (Fig. 2F), measured from the class averages of autoinhibited KIF5A molecules. This is equivalent to ∼207 residues of coiled coil, assuming a rise per residue of 0.15 nm. The measured length is slightly shorter than the expected length (combined length of CC1 and 2, ∼36.3 nm, 242 residues) assuming the bend occurs in the elbow (Fig. 1B) as observed for KIF5B and C. This suggests that the organisation of the coiled coils into the KIF5A autoinhibited structure is more complex than a simple bend at the elbow. The proline residue (P687) in the elbow sequence, is expected to disrupt the coiled coil and could facilitate bending. A proline residue at this position is conserved in KIF5A across a wide range of mammalian species, although absent in KIF5C and KIF5B (Fig. 1B).

The distance between the central regions of the two motor domains (Fig. 2G; 6.6 ± 1.1 nm) is consistent with the known distance between the two motor domains in the autoinhibited structure for *Drosophila* kinesin-1 motor domains crosslinked by the IAK motif (5.6 nm: PDB 2Y65) (16). The distance between the motor domains and the additional density (bright region of negative stain, or ‘bulge’) about halfway along the stalk was 16.7± 3.6 nm (Fig. 2H), equivalent to ∼111 residues of coiled coil. A bulge in a similar position was seen for KIF5B (19) and a complex of KLC1 and rat KIF5C (18), in which a predicted six-helix bundle is formed at the interface between these two proteins. Even though there are no light chains present in our molecules, it is likely that this bulge corresponds to hinge-2 and the KLC binding interface, which should be juxtaposed at this point. Hinge-2 and the KLC interface both contain intrinsically disordered regions (∼20 residues for hinge-2) and presumably collect additional stain that results in the appearance of a bulge in the nsEM images of KIF5A.

A second bulge, closer to the motor domain (termed the shoulder), reported for the complex of rat KIF5C and KLC1-TPR (tetratricopeptide repeat) domains (18) is absent. The complex between rat KIF5C and a mutant KLC1 that lacks TPR domains also lacked this bulge. This bulge was suggested to be a second KLC1-KIF5C interaction site (18). Thus, the absence of a shoulder in our images is consistent with the absence of light chains in the purified KIF5A.

### Molecular architecture of autoinhibited KIF5A

To better understand the nsEM images, we developed a model for the overall structure of autoinhibited KIF5A, using a combination of Alphafold2 (29, 30), 3D-reconstruction of the nsEM images and crosslinking mass-spectrometry (XL-MS) (Fig. 3). An Alphafold2 model of autoinhibited KIF5A was generated from 4 initial overlapping regions of KIF5A (Fig. 3A), motor-CC1 (aa 1-540), CC1-CC2 (aa 401-690), CC3 (aa 691-820) and CC4 and C-terminal tail (aa 821-1034). Interestingly, the initial Alphafold2 model of the motor, neck coil and CC1, positions CC1 between the two motor domains and the neck coil protrudes below (Fig. 3A). We noticed that this was somewhat consistent with a region of additional density in the 3D nsEM model below the two motor domains (Fig. 3B). This protrusion was also evident in some class averages (Fig. 3D). The 4 segments were positioned loosely in the 3D nsEM density map (Fig. 3B) and refined to generate a reasonable model of full length regulated KIF5A. This was further refined iteratively and validated, using additional information from XL-MS (Fig. 4) to generate the final model (Fig 3C). The final model also reconciles the structure of the autoinhibited motors, from the original crystal structure *of Drosophila* KHC (16) compared to their position in the starting model (Fig. 3A).

**Fig. 3.**
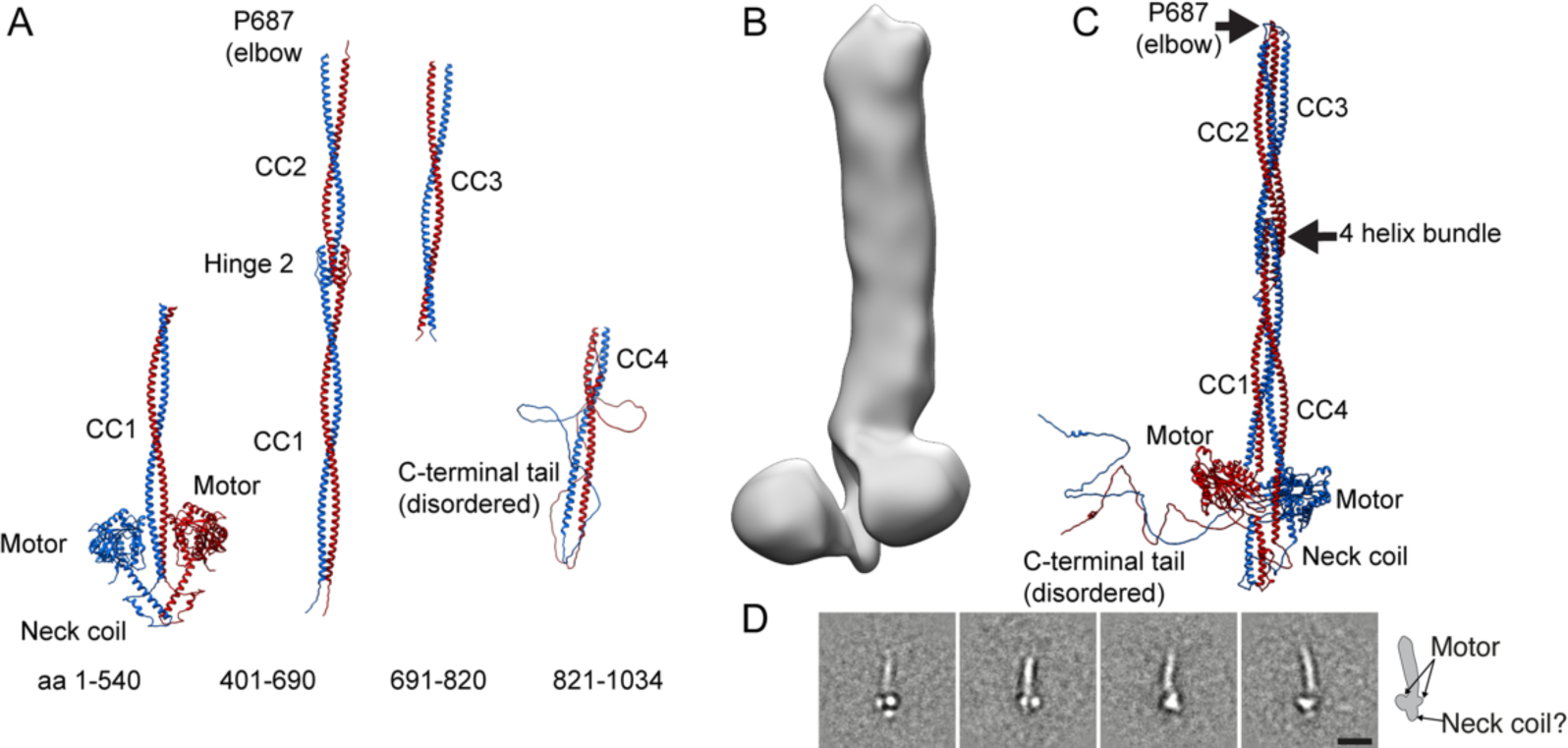
Alphafold model of autoinhibited KIF5A: **A:** Alphafold models of segments of KIF5A used to build the final structure in **C**. **B:** 3D nsEM image of KIF5A. **C:** Final model of the autoinhibited KIF5A. **D**: Class averages with apparent neck domain bulge protruding from below the motor domains (Shown in inset). Scale bar 20 nm.

**Fig. 4:**
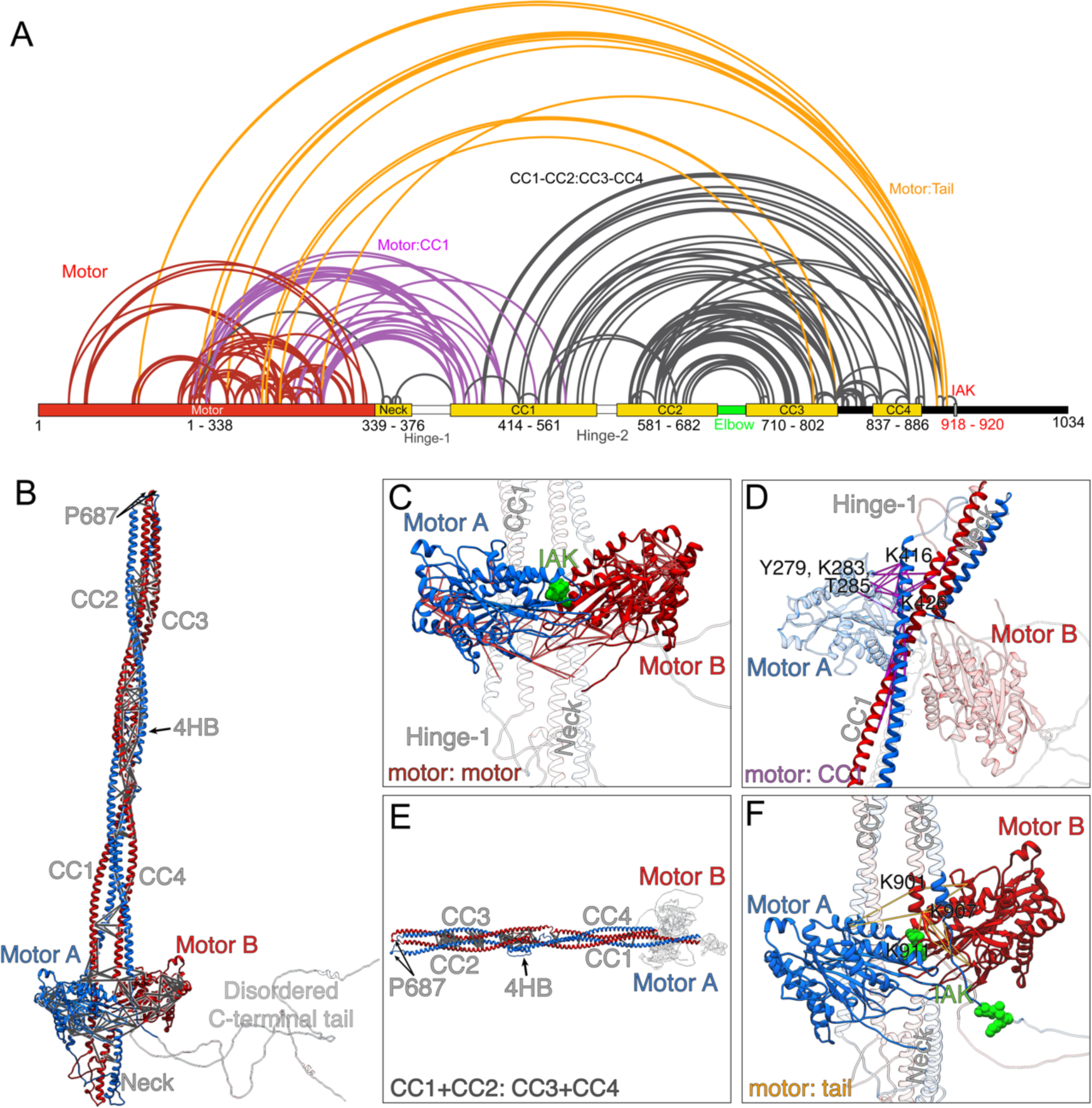
Crosslinking mass spectrometry data for autoinhibited KIF5A. **A:** Crosslinked residues are indicated by the lines drawn. Red lines indicate crosslinks formed within the motor domains, purple indicates crosslinks between the motor and CC1, orange lines indicate cross links between the proximal region of the disordered C-terminal tail, close to the IAK motif and the motor domain, and black lines indicate crosslinks between CC1, CC2, CC3 and CC4. **B:** shows the crosslinks super-imposed on the full-length model. **C**: shows crosslinks present in the motor domains (Red lines in A), with IAK motif from the tail shown in green. The minimum and maximum detected crosslink lengths were 5.8 Å and 26.7 Å, respectively. **D:** shows crosslinks between the motor and CC1 (purple lines in A). The minimum and maximum detected crosslink lengths detected were 6.7 Å and 25.4 Å, respectively. **E.** Shows crosslinks between coiled coils (black lines in A). The minimum and maximum detected crosslinks detected were 9.3 Å and 27.0 Å, respectively. **F.** shows crosslinks between the proximal region of the C-terminal and the motor (orange lines in A). The minimum and maximum detected crosslink lengths were 11.5 Å and 26.5 Å, respectively.

In the final model, bending of KIF5A at P687, together with the positioning of each of the coiled-coil domains allows the IAK motif in the tail region of KIF5A to be positioned close to and interact with the motor domains, in the autoinhibited molecule (Fig. 3C). The distance between the centre of mass between the motor domains and P687 is 32 nm, which matches measurements from the negative stain data. The neck coil is positioned below the motor domains and the proximal region of CC1 lies between them. The interaction between CC1 and CC2 in the region of hinge-2 generates a 4-helix bundle as indicated (Fig. 3C), similar to that previously reported for KIF5B and KIF5C (18, 19). The position of this bundle in our model (∼17.8 nm away from the motor domains) is consistent with the position of the bulge seen in our 2D class averages (Fig. 2).

Crosslinking mass spectrometry (XL-MS) data showed extensive crosslinks between residues in the motor domains, between the motor and CC1, between the coiled coils, and between the motor and the tail (Fig. 4A, Supplemental Table 2). Extensive crosslinks between residues within and between the motor domains are consistent with known distances in the *Drosophila* KHC crystal structure of the autoinhibited motor and IAK peptide (16), suggesting that the XL-MS data reports crosslinked residues faithfully (Fig. 4A-C). The measured distances for 38 out of 54 crosslinked pairs in this region were within the theoretical limit of the BS3 crosslinker distance constraints of up to ∼27Å with a tolerance of an extra 3 Å (31).

To position the neck coil and CC1 as shown in the final Alphafold2 model and be consistent with the crosslinking data, the molecule must fold back at hinge-1 to enable the motor domains to interact with CC1 (Fig. 4 A,B and D). Extensive crosslinks between residues in one of the two motor domains (MotorA, blue) and residues in CC1 are comprised of Loop12, the α5 helix (G272 – L292) and the phosphate binding loop (K257) in MotorA and K416, K426, & T417 in CC1 (Fig. 4C), further support our final Alphafold2 model (Fig. 3C).

The presence of multiple crosslinks between CC1 and CC2, and between CC3 and CC4, suggest that the autoinhibited structure is also likely stabilised by interactions between these regions of coiled coil (Fig. 4A,B and E). The high density and positioning of crosslinks between CC2 and CC3 is consistent with folding of the molecule at the elbow (P687) and places CC2 and CC3 in apposition.

The proximal region of the C-terminal tail (residues 890-910) close to the IAK motif is crosslinked to the motor domain (Fig. 4.A,B and F). This is consistent with the known interaction of the IAK motif from one of the two C-terminal tails with the motor domain, as shown by the crystal structure of the motor & IAK peptide complex (16). For example, residues K901, 907, 911 in the auxiliary MT binding site in the C-terminal tail (Fig. 1B) that lie upstream of the IAK motif, make numerous crosslinks with residues in loop 5 and the α3 helix (S176 – R191) of both motor domains.

We did not detect any crosslinks between the remaining downstream sequence of the C-terminal tail (112 residues) and the rest of the molecule. We did detect dead-end modifications on lysine, serine, tyrosine and threonine residues (35 out of 112 residues) within this C-terminal sequence suggesting that they are accessible and could have formed crosslinks with other regions of KIF5A, if they were involved in any interactions. It is possible that this region of the molecule, likely to be intrinsically disordered and highly dynamic, does not interact with other regions of the molecule, and is not directly involved in stabilising the autoinhibited state. However, we cannot rule out that a different crosslinker, with a longer spacer arm than BS3 used here (11 Å) such as BS(PEG)5 (21.7 Å) or even BS(PEG)9 (35.8 Å) may have shown some crosslinks.

### Mapping mutations in autoinhibited KIF5A

Of the three kinesin-1 isoforms (KIF5A, KIF5B and KIF5C), mutations in KIF5A are most common, with >80 missense mutations and 15 splicing mutations reported (Supplemental Table 1, Fig. 1B). In contrast, just over 20 mutations have been reported for KIF5C, which predominantly result in neurodevelopmental disorders, such as cortical development and microcephaly (32, 33), likely through a key role in neuronal polarisation (34). Very few mutations (less than 5) have been reported for KIF5B.

Missense mutations in KIF5A are found throughout the sequence (Fig. 5A-B). However, mutations in the motor domain are more commonly associated with spastic paraplegia, while mutations in the tail are more commonly associated with amyotrophic lateral sclerosis (Fig. 5C). About 50% (43 mutations) occur in the motor domain (Fig. 1B, Fig 5A,B). Most of these are in the phosphate binding loop and the α4 helix, switches 1 and 2 (3 out of 6 residues are mutated in each: Fig. 1B). Switch 1 and 2 and the phosphate binding loop are critical in co-ordinating nucleotide binding, and communicating nucleotide state to the motor, to drive kinesin motility. Switch 1 mutations reduce or abolish kinesin motility in vitro (35). None of the residues implicated in the binding of the IAK motif in the tail to the motor domain are mutated, although some nearby residues are, and these are all implicated in Charcot Marie Tooth Disease (CTMD) (Fig. 1B, Supplemental Table 1).

**Fig. 5.**
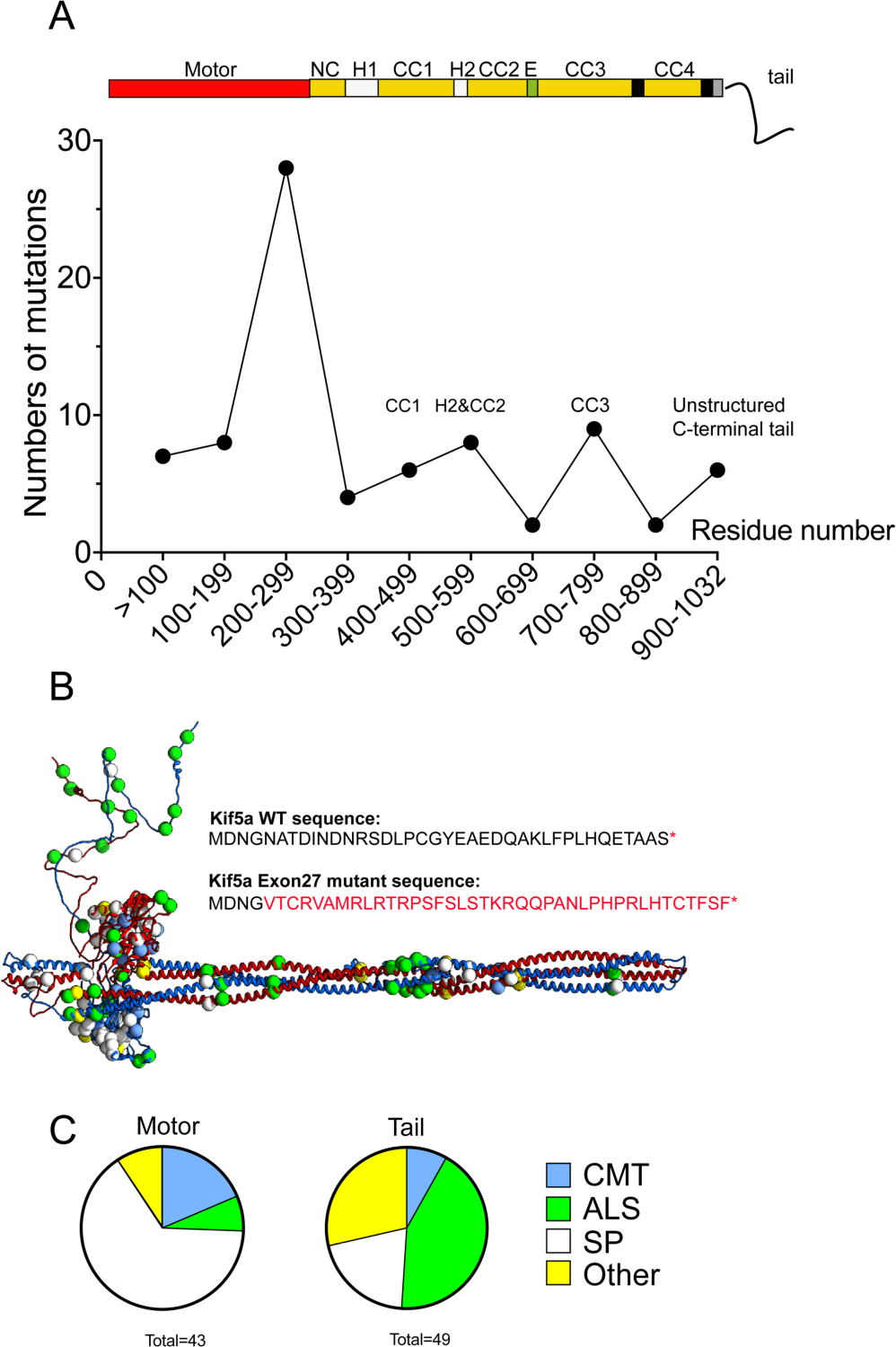
The location of disease-causing mutations in KIF5A. A: shows the numbers of missense mutations in KIF5A calculated for every 100 residues and plotted as shown. A diagram of KIF5A domains is shown above the plot for context. B: mutations in KIF5A plotted onto the model structure are shown as green circles. The altered sequence that results from exon 27 skipping is also shown. C: Further analysis of the KIF5A mutations that cause different diseases: CMT (Charcot Marie Tooth), ALS (Amyotrophic lateral sclerosis), SP (Spastic Paraplegia type 10) and others shown separately as a Pie Chart for the motor and the tail. SP mutations are most commonly found in the motor, while ALS causing mutations are most commonly found in the tail.

In the coiled-coil region of KIF5A, seven missense mutations are found in CC1, four in CC2, nine in CC3 and three in CC4, with three mutations in hinge-2 (Fig. 1B, 5A). Thus, mutations in CC3 are the most common in the tail region (Supplemental Table 1). If they destabilise the coiled coil, they have the potential to destabilise the autoinhibited molecules. Mutations in and around hinge-2 could also destabilise the autoinhibited state, and/or interaction with light chains. Plotting the mutations onto the model structure demonstrates the large number of ALS mutations found close to the position of the 4-helix bundle, in the central region of the shaft (Fig. 5B) again raising the possibility that these mutations could affect the stability of the autoinhibited state.

The potential effect of mutations in the C-terminal sequence are much less clear. Eight missense mutations are found in the C-terminal sequence unique to KIF5A, a disordered region that does not appear to interact with any other regions of the molecule in the autoinhibited state, and thus would not be expected to affect the stability of the autoinhibited state. Many of the splicing mutations result in the skipping of exon 27, which does not remove the IAK motif, but alters the C-terminal sequence after residue 998. Exon 27 mutants have been reported to be constitutively active (23, 25), but also to form aggregates (24–26) and can fail to transport mitochondria correctly (26). The increased number of negatively charged residues in the substituted tail sequence has been suggested to interfere with the ability of the positively charged IAK motif to bind to the motor domains thereby resulting in a constitutively active molecule. However, this region of substituted sequence lies within the intrinsically disordered region that does not appear to interact with the motor domains in the autoinhibited molecule. It is possible that aggregation of KIF5A exon 27 mutant indirectly interferes with its ability to associate with TRAK2.

To better understand how the exon 27 mutant affects the autoinhibited state, we expressed and purified full length KIF5A comprising the exon 27 mutant sequence (Fig. 5B). Mass photometry showed a similar distribution of molecular weights to wild type (WT) (Supplemental Fig. 2A). This mutant was also able to adopt the autoinhibited state, as we could observe autoinhibited molecules, prepared in the same way as for wild type KIF5A, using negative-stain EM experiments (Supplemental Fig. 2B).

Surprisingly, XL-MS showed an increased number of crosslinks for the autoinhibited exon 27 mutant KIF5A compared to those for WT KIF5A (Supplemental Fig. 2C, Supplemental Table 1). There was often an increased number of crosslinks in same residue for the exon 27 mutant, compared to WT. For example, in WT KIF5A, K99 in the motor domain (loop 5) for both motor A and motor B (chain A, chain B) is crosslinked to K188 and S189 in the same motor domain, and to K901 and K907 in chain B (10 residues downstream of the IAK motif) in the tail. In the exon 27 mutant, in addition to these crosslinks, crosslinks between K99 and residues T196, S202 and S203 (α3a region of the motor domain) were also present. Similarly, in wild type KIF5A, K167 in motors A and B is crosslinked to S155 and K283 in the same motor domain, K167 in motor A is crosslinked to residues K416 and K426 in chain A (proximal region of CC1), and K167 in motor B is crosslinked to residues K907 and K911 in chain B. In the exon 27 mutant, the crosslink between K167 and S155 is absent in both motors, K167 in both motors is crosslinked to K416 and K426 in chain A and B, respectively,and K167 in motor B is crosslinked to residues 907 and 911 in chain B, and additionally to Y927 in chain B (6 residues distal to the IAK motif), whereas K167 in chain A is additionally crosslinked to K920 in chain B, the K residue in the IAK motif. One possible interpretation of these results is that the autoinhibited state for the exon 27 mutant is more labile that WT, which would allow specific residues to form more crosslinks. However, we might expect numbers of crosslinks to be reduced in the exon 27 mutant if the autoinhibited state was significantly disrupted.

Finally, the exon 27 KIF5A mutant expressed as a GFP-fusion protein in mammalian cells, showed aggregates, as previously reported (23) whereas wild type protein had a diffuse organisation (Fig. 6). GFP-KIF5A constructs with one of three missense mutations in the C-terminal region (P896L, D1002G or R1007G) showed a similar pattern to wild type KIF5A. This suggests that exon 27 KIF5A mutant affects autoinhibition, whereas missense mutations, in the disordered region of the C-terminal tail do not.

**Fig. 6.**
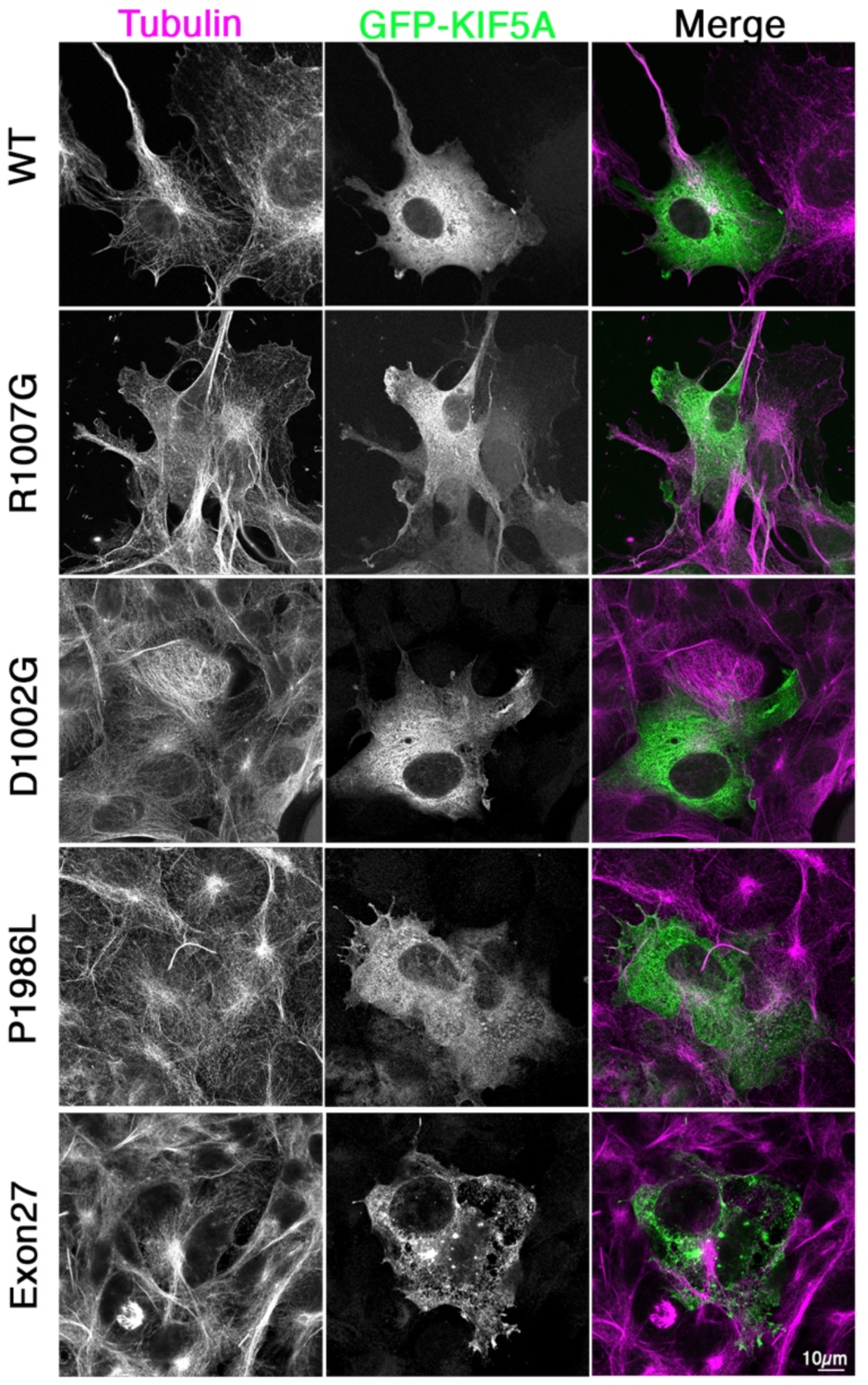
Airyscan confocal images of GFP-tagged KIF5A WT and mutants in cells. The images show. COS7 cells transfected with WT and mutant GFP-KIF5A constructs as indicated (shown in green in the merged image), and co-stained for tubulin (shown in red in the merged image) and DAPI (blue in the merged image). Scale bar 10 µm.

## Discussion

Here, using a combination of nsEM, Alphafold modelling and XL-MS, we determined a model structure for autoinhibited KIF5A. Plotting KIF5A disease mutations onto this structure demonstrate that many mutations within the coiled-coil tail appear to be in key regions of the molecule that help to stabilise the autoinhibited structure. The additional sequence in the C-terminal region of KIF5A, not found in the other kinesin-1 isoforms, is likely to be intrinsically disordered and not directly involved in forming the autoinhibited state. However, the change to the C-terminal sequence that results from skipping exon 27 does appear to destabilise the autoinhibited state as evidenced by changes to the XL-MS data and to the GFP-KIF5A staining pattern in cells, while single missense mutations in the C-terminal tail do not.

The model structure we obtained for the KIF5A heavy chain is consistent with the previous model structures for KIF5B and KIF5C (18, 19). It shows that KIF5A does not simply fold in half via hinge-2 between CC1 and CC2 and previously suggested (12, 36). Instead, the main bend that allows KIF5A to form the autoinhibited state is in the elbow region, which in KIF5A contains a proline residue (P687) likely to locally disrupt the coiled coil, and thus act as a point of flexibility to enable formation of the autoinhibited state. Hinge-2 is involved in the formation of an antiparallel 4-helix bundles along the shaft of the autoinhibited molecule, and has been shown to act as the interface for KLC binding in autoinhibited molecules, in which KLC binding stabilises the autoinhibited state (18, 19).

The autoinhibited state formed by the heavy chain in the absence of light chains is likely stabilised by multiple interactions between the motor and the C-terminal tail, the motor and CC1-CC2 and CC3-CC4, and interactions between the two motors. Flexibility arising from hinge-1 enables the interaction of CC1 with motorA in the autoinhibited state, placing the coiled coil close to the phosphate binding loop in the motor and blocking the microtubule-binding interface of the motor domain. It also enables the positioning of the neck domains at an angle of 180° to the stalk domain. The positioning of CC1 across the surface of motorA brings the distal region of CC4 and the IAK motif close to the two motors, enabling the IAK motif from one of the two chains to interact with the two motors in the autoinhibited state, consistent with previous observations.

The skipping of exon 27 in KIF5A, in which the last 34 residues in the C-terminal tail are replaced by a novel 39 amino acid sequence, has previously been suggested to autoactivate KIF5A. Additional experiments showed that this mutation conferred a toxic gain of function and aggregation of KIF5A and increased motility for the exon 27 mutant in vitro compared to wild type (23, 25), but it has also been shown to disrupt mitochondrial transport in neurones (26). In partial agreement, our data also shows that the exon 27 mutant is aggregated in cells, but it is less clear if the mutant forms a more open state as evidence from the XL-MS and nsEM data. The exon 27 mutant can still be crosslinked into the same autoinhibited conformation as wild type KIF5A. It is possible that the increased number of crosslinks that we observed in the exon 27 mutant could be interpreted as arising from a greater instability of the autoinhibited state, allowing specific residues to interact more widely. However, adding TRAK1 to KIF5B, which destabilises the autoinhibited state markedly reduced the number of crosslinks (19). It is possible that the substituted C-terminal sequence is more likely to promote self-aggregation in the exon-27 mutant, which in turn destabilises the autoinhibited state and promotes auto-activation as reported.

We found no evidence for any crosslinks between the last ∼70 residues of the C-terminal tail of KIF5A and the motor and/or shaft of the autoinhibited kinesin molecule (Fig. 4). We cannot rule out that a crosslinker with a longer spacer arm may have revealed some crosslinks. However, the presence of an extended C-terminal tail is not required for formation of an autoinhibited state in KIF5B and KIF5C, where this extended C-terminal sequence is absent. The specific role of the intrinsically disordered C-terminal tail in the regulation of KIF5A thus remains unclear. It is likely to be important, as it contains multiple missense mutations that have been linked to disease (Fig. 1B, Supplemental Table 1). However, the 3 missense mutations we tested do not appear to activate KIF5A when expressed in cells, as no aggregates were detected. Intrinsically disordered regions, such as that found in the C-terminal tail of KIF5A, can be involved in protein-protein interactions and mutations within them have been implicated in disease (37). Thus, mutations in the C-terminal tail of KIF5A, may possibly have a stronger effect of protein-protein interactions than on regulation.

Overall, this work shows a general agreement in the mechanism by which each of the three kinesin-1 isoforms fold up into an autoinhibited state as a homodimer. The formation of an autoinhibited state may facilitate the transport or diffusion of the motors to sites of action, as previously suggested (38, 39). However, it is not yet completely clear if kinesin-1 is able to form autoinhibited homodimers, autoinhibited heterotetramers (in complex with light chains or other adaptor proteins) or both in cells. Moreover, kinesin-1 molecules may also fold up into an autoinhibited state while directly associated with cargo and/or be released into the cytoplasm (40). Finally, the findings presented here have broad relevance to disease mutations in KIF5A, and it will be interesting to see in future work if specific mutations destabilise the autoinhibited state and/or affect recruit to cargo.

### Experimental Procedures

#### Sequence alignment

The amino acid sequences for KIF5A (Q12840),KIF5B (P33176) and KIF5C (O60280), obtained from UNIPROT, were imported into JALVIEW (41) and aligned using Muscle. The resulting alignment was exported and annotated in Illustrator (Adobe) to highlight the different domains. Coiled coil analysis was performed using Waggawagga ((42) available online at https://waggawagga.motorprotein.de/). The Marcoil prediction output was selected and manually checked to generate the coiled coil annotations in Fig. 1B. Single amino acid mutations in KIF5A listed by the Human Genome Mapping Database (HGMD: accessed December 2022) were added to the figure. Mutations are listed in Supplemental Table 1, together with associated references. There were 81 missense mutations and 15 splicing mutations, most of which affect splicing of exon 28.

#### Protein Expression, Purification, and Reagents

The coding sequence for human KIF5A (UNIPROT Q12840: Genscript clone ID: OHu18844: NM_004984.4) and for the exon-27 splice mutant, with a C-terminal FLAG tag (sequence: DYKDDDDK) immediately following the C-terminal serine residue were synthesised and subcloned into the pFastBac1 vector (Invitrogen) by Genscript. Creation and amplification of recombinant baculoviruses, expression and purification of proteins were as described previously (43). Briefly, an MOI of 5 for the KIF5A virus was used to infect the *Sf*9 cells for protein expression. Baculovirus infected *Sf*9 cells were grown for 72 hr and harvested by sedimentation. Cell pellets were stored at −80°C.

To purify the protein, frozen pellets were thawed and homogenized on ice using a ground glass homogenizer in buffer A (10 mM MOPS (pH 7.4), 5 mM MgCl2, 0.1 mM EGTA) supplemented with 0.5 M NaCl, 2 mM ATP, 0.1 mM phenylmethylsulfonyl fluoride and protease inhibitor cocktail (Roche). The proteins were purified by FLAG-affinity chromatography using M2 FLAG affinity gel (Sigma-Aldrich) and eluted in buffer A supplemented with 0.5 M NaCl, and 0.5 mg/ml of FLAG peptide (Sigma). The eluted proteins were dialysed overnight in buffer A supplemented with 0.5 M NaCl and 1 mM DTT. Protein concentration was determined using a nanodrop, with the calculated extinction coefficient (ε=57190; 114380 for a dimer). The protein was flash frozen in liquid nitrogen in ∼ 20 µl aliquots and stored in liquid nitrogen until used. Approximately 5 mg of protein was obtained from 0.75×10^9^ cells.

For GFP-KIF5A, the same sequences were subcloned into pEGFP-C1, to position EGFP at the N-terminus as we did previously for KIF5C (44). In addition, 3 additional missense mutations were generated by Genscript: P896L, D1002G, and R1007G for GFP-KIF5A. All GFP-KIF5A constructs were maxiprepped and transfected into COS-7 cells (ATCC) using Fugene6 (3:1 Fugene:DNA ratio). COS-7 cells (mycoplasma free) were cultured as described (44), and cells were plated onto detergent-washed 13mm diameter glass coverslips (# 1.5) at least 8 hours prior to transfection. Cells were fixed using 4% paraformaldehyde in phosphate buffered saline (PBS), co-stained for tubulin using an Affimer specific for tubulin (45) conjugated to Alexa 647 and imaged by Airyscan confocal microscopy.

#### Electron Microscopy and Image Processing

KIF5A was diluted into low salt buffer (150mM KCl, 10mM MOPS (pH 7.2), 2mM MgCl_2_, 1mM EGTA, 1mM ATP) to a concentration of 2 µM. The sample was then crosslinked by adding BS3 (bis(sulfosuccinimidyl)suberate: Thermo Fisher Scientific) to a final concentration of 1 mM, incubated at 25°C for 30 min and quenched using 100mM Tris-HCl, pH 8.0. Crosslinked KIF5A was diluted to a concentration of 20nM in low salt buffer. This protein was applied to glow discharged, continuous carbon-coated grids and negatively stained with 1% uranyl acetate. Images were captured at x50,000 magnification on a FEI CETA CCD camera using a FEI Technai F20 (ThermoFisher) transmission electron microscope operating at 120 kV and digitized at 0.2 nm/pixel.

Particles were picked using Relion3.1 (46) and image processing was performed using IMAGIC5 (Image sciences)(47). The particle stack contained 4985 particles. K-means classification was performed by using masks based on the variance of the global average. Class averages were then analysed in ImageJ.

The resulting particle stack was exported to cryosparc (48) and subjected to ab initio reconstruction followed by non-uniform refinement. The resulting map was 24 angstrom resolution.

#### Single molecule Mass Photometry

Single-molecule landing assays, data acquisition and image processing were performed as previously described (49, 50) using the Refeyn One^MP^ mass photometer. Briefly, microscope coverslips (#1.5, 24 × 50 mm and 24 × 14 mm, Fisher Scientific) were cleaned consecutively with isopropanol and water then dried using a stream of clean nitrogen. A silicon gasket was applied onto the cleaned coverslip and gentle pressure was applied to ensure it was firmly stuck. PBS was applied to the cleaned coverslip to buffer-blank before 25nM of native KIF5A was applied to the coverslip. Data was collected in a ∼3 μm × 10 μm field of view at an acquisition rate of 1 kHz for 120 s. All measurements were carried out at room temperature (∼23 °C). Images were processed using the manufacturer’s software (Refeyn, UK). The conversions between molecular mass and interferometric contrast were calibrated using protein standards of known molecular weight in PBS. The histograms generated from the measurements were fit with a Gaussian using Prism (Graph-Pad). The positions of the peaks for each Gaussian were used to determine the mass of the kinesin molecules, using the plot generated from the protein standards.

#### Crosslinking mass spectrometry

2μM purified KIF5A was prepared in 30 μl low salt buffer to allow the shutdown state to form. 2 mg of BS3 (Thermo Fisher Scientific), which mainly targets lysines, but will also crosslink serine, threonine and tyrosine (51)was dissolved in 117 μl low salt buffer to a final concentration of 30 mM. 1 µM BS3 was added to the shutdown KIF5A in solution. The reaction mixtures were incubated at 25°C for 30 min, after which they were quenched with Tris-HCl (pH 8) at a final concentration of 100 mM. A 5 μl sample was used for SDS-PAGE to evaluate the crosslinking quality for each reaction.

Following the crosslinking reaction, samples were mixed with 10% SDS in equal volumes and then subjected to S-TRAP^TM^ digestion, as per instructions (PROTIFI, NY, USA). Briefly, Samples were initially reduced and alkylated using 20 mM DTT for 10 minutes and 40 mM IAA for 30 minutes respectively followed by acidification of samples using 5.5% phosphoric acid. Acidified samples were then trapped on S-TRAP columns after the addition of sample buffer (100 mM triethylammonium bicarbonate buffer (TEAB) in 90% methanol) and 1 µg trypsin. The column was then washed with sample buffer, and the column was incubated at 47°C for 90 minutes. Eluted peptides were concentrated in a speedvac concentrator and reconstituted in 0.1% formic acid (FA).

LC-MS/MS (LC: Liquid Chromatography, MS: mass spectrometry) analyses of crosslinked peptides were performed on an Orbitrap Eclipse Tribrid mass spectrometer (Thermo Fisher Scientific) coupled to a Vanquish Neo UHPLC system (Thermo Fisher Scientific). Prior to LC separation, tryptic digests were online concentrated and desalted using a trapping column (300 μm × 5 mm, μPrecolumn, 5μm particles, Acclaim PepMap100 C18, Thermo Fisher Scientific) at room temperature. After washing of the trapping column with 0.1 % FA, the peptides were eluted (flow rate – 0.25 nl/min) from the trapping column onto an analytical column (EASY spray column, Acclaim Pepmap100 C18, 2µm particles, 75 μm × 500 mm, Thermo Fisher Scientific) at 45 °C by approximately 95 min linear gradient program (2-50 % of mobile phase B; mobile phase A: 0.1 % FA in water; mobile phase B: 0.1 % FA in 80 % ACN). Equilibration of the trapping column and the analytical column was done prior to sample injection to the sample loop. The analytical column with the emitter was directly connected to the ion source.

MS data was acquired in a data-dependent strategy (DDA). The mass spectrometric settings for MS1 scans used were resolution set to 120,000, AGC of 3 × 106, maximum injection time of 50 ms, scanning from 380–1450 m/z in profile mode. With z = 3–8 and using isolation window of 1.4 m/z. Fragmentation done by HCD using stepped normalized collision energies (30 ± 6). Fragment ion scans were acquired at a resolution of 60,000, AGC of 5 × 104, maximum injection time of 120 ms. Dynamic exclusion was enabled for 30 s (including isotopes). Each LC-MS run took 97 min. We repeated these experiments 5 times for the wild type KIF5A, and 3 times for the exon-27 mutant KIF5A.

The mass spectrometric RAW data files were analyzed using the XlinkX, Proteome Discoverer software (Thermo Scientific, version 3.0.0757). and Merox (2.0.1.4) software. Searches were performed against a FASTA file containing proteins of interest. Oxidation of methionine, deamidation (N, Q) as optional modification and carbamidomethyl on cysteine as static modification were used. Trypsin (full) enzyme with 2 allowed miss-cleavages were set. Mass spectra were searched using precursor ion tolerance 5 ppm and fragment ion tolerance 10 ppm. Peptides and proteins FDR (false discovery rate) threshold was set to <0.01 and the cut off score was set to 50. In the data presented here (Supplemental Table 2), the crosslinks identified appeared in all 5 runs for wild type KIF5A, and in all 3 runs for exon-27 mutant KIF5A and are within the cut-off distance of ∼27Å.

#### Alphafold structure prediction of kinesin-1 and fitting

The structure of the human autoinhibited KIF5A motor domains (1–345) was generated by substituting the amino acid sequence of 2Y65 (*Drosophila Melanogaster* kinesin-1 motor domain dimer-tail complex crystal structure (16): amino acids 8-351) with that of human KIF5A (Uniprot Q12840) using Modeller9.23 (52). This yielded a homology model of the autoinhibited motor domains; termed autoinhibited KIF5A motors (aa1-345) from here onwards.

Models of four fragments of KIF5A; motor-CC1 (aa 1-540; composed of the motor domains and neck and beginning of CC1), CC1-CC2 (aa 401-690), CC3 (aa 691-820) and CC4 and C-terminal tail (aa 821-1034) domains were generated using AlphaFold Multimer colabfold (53) using no existing templates and with 3 recycles. The C-terminal region of each model fragment generated corresponds to regions predicted to have a low confidence score of forming a coiled coil as predicted by MARCOIL (54). These disordered regions provide extra degrees of freedom to fit the coiled-coil model within the 3DEM density map (discussed below). The top ranked model, based on confidence scores from each model fragment was taken forward for fitting in the 3D density map.

The neck coil of the autoinhibited KIF5A motors (aa1-345) model was extended downwards to R373 and joined to the motor-CC1 Alphafold model at N374; with the overlapping sequence (1-373aa) from the motor-CC1 model being deleted. This yielded the autoinhibited KIF5A motor-CC1 (1-540aa) model.

CC1 was manually positioned to be in close proximity to the motor domains as shown by the motor-CC1 (1–540) Alphafold model (Fig. 3A). The residues from the CC1-CC2 model (401–540) were superposed onto CC1 (401–540) from the autoinhibited KIF5A motor-CC1 (1-540aa) model created above. Overlapping sequences were deleted and the models were joined to yield the autoinhibited KIF5A motor-CC1 and 2 (1-690aa) model. The autoinhibited KIF5A motor-CC1 and 2 (1-690aa), CC3, and CC4 and C-terminal tail models were positioned loosely in the 3DEM density map, using the flexibility provided by the disordered regions at the N-& C-terminus of each model to improve the fit within the map. Additionally, using the additional positional restraints provided by the crosslinking mass spectrometry to iteratively improve the fit of these three discrete segments with the 3D density map. Once satisfied with the fit, the segments were joined. Coot (55) was used to correct changes, such as long bonds introduced between coiled coil segments as a result of the manual fitting procedure.

## Supporting Information

This article includes Supporting Information.

## Acknowledgements

We thank the staff in the Mass Spectrometry Facility for help and advice.

## Funding and additional information

This work was supported by a Wellcome Trust Investigator award to MP: WT 223125/Z/21/Z. The Airyscan confocal Microscope was funded by the Wellcome Trust: WT104918MA. Uzrama Fatima was a MSc Student in Biosciences at the UoL. Ines Caramujo was an MBiol. Student at the University of Leeds. The eclipse mass-spectrometer was funded by the Wellcome Trust : 223810/Z/21/Z.

## Data availability

The XL-MS data is provided in Supplemental Table 2 in Supporting Information.

