## Supplemental Data for "A Multiscale approach reveals the molecular architecture of the autoinhibited kinesin KIF5A"

Supplemental Figure 1
Supplemental Figure 2
Supplemental Table 1
Supplemental Table 2

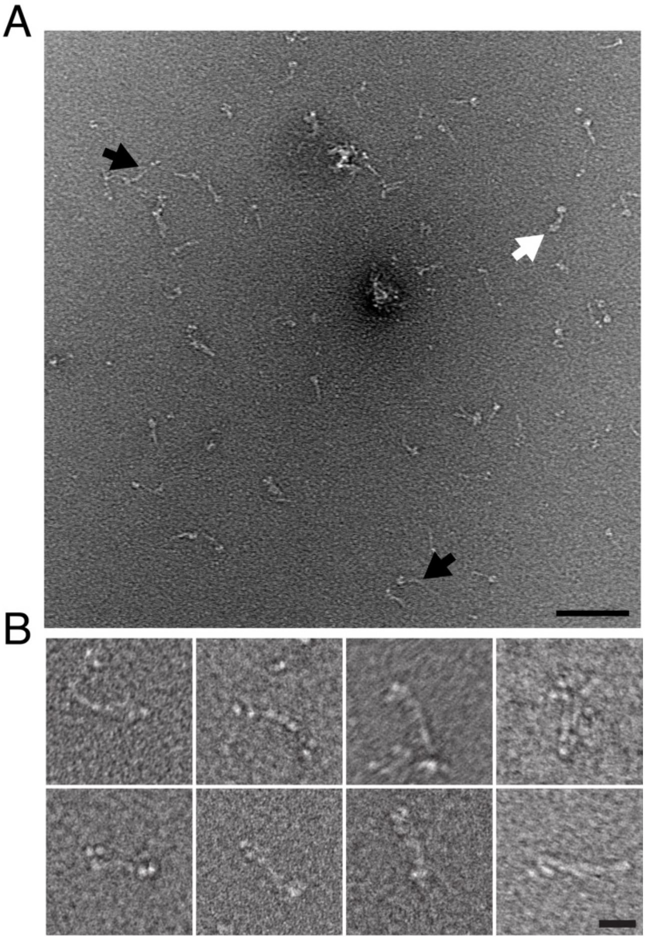


**Supplemental Fig. 1.** NsEM images of WT KIF5A show the presence of occasional tetramers and open heads. **A:** Representative field of view. Black arrows indicate open heads White arrow indicates a tetramer. Scale bar: 100nm **B:** Montage of individual images of tetramers. Scale bar : 20 nm

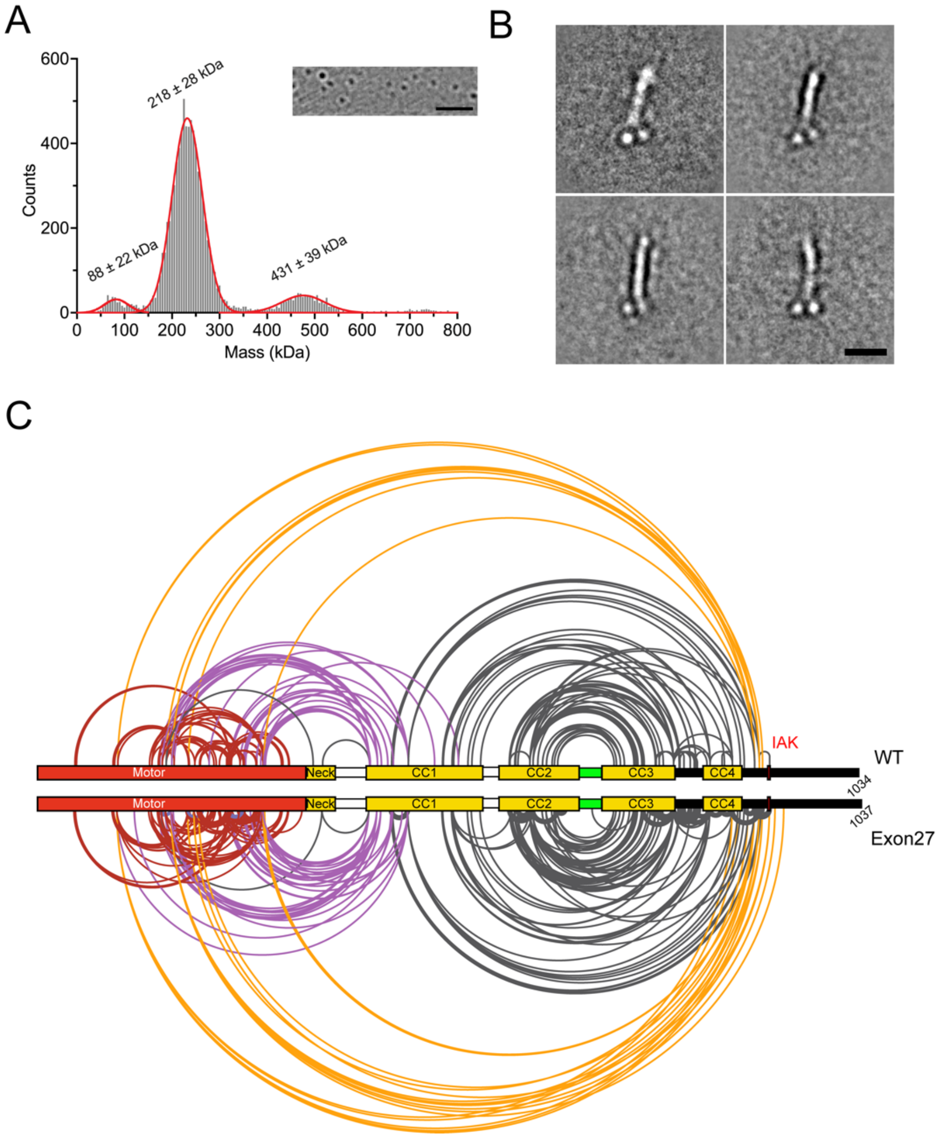


**Supplemental Fig. 2. Analysis of exon 27 KIF5A mutant. A:** mass spectrometry data for the mutant KIF5A. Numbers above each peak show the mean molecular mass for 3 measurements ± S.D. The image insert shows the type of image obtained in this experiment. Scale bar 2 μm: **B**: Montage of negative stain EM class averages mutant KIF5A: scale bar 20 nm. **C**: XL-MS data for the exon 27 mutant, compared to the data presented in Fig. 4 in the main paper for the WT KIF5A. Cross-linked residues are indicated by the lines drawn. Red lines indicate crosslinks formed within the motor domains, purple indicates cross links between the motor and CC1, orange lines indicate cross links between the proximal region of the disordered C-terminal tail, close to the IAK motif and the motor domain, and black lines indicate cross links between CC1, CC2, CC3 and CC4.

**Supplemental Table 1: Mutations in KIF5A**

| Mutation | Kinesin region | Disease | Reference |
|---|---|---|---|
| V12A | Motor | Charcot-Marie-Tooth disease, type 2 | (1, 2) |
| K29R | Motor | Amyotrophic lateral sclerosis | (1) |
| P24S | Motor | Amyotrophic lateral sclerosis | (1) |
| Y63C | Motor | Spastic Paraplegia | (3) |
| D73N | Motor | Charcot-Marie-Tooth disease, type 2 | (1) |
| V74A | Motor | Amyotrophic lateral sclerosis | (4) |
| Q87E | Motor | Spastic Paraplegia | (5) |
| R111P | Motor | Charcot-Marie-Tooth disease, type 2 | (6) |
| K132R | Motor | Charcot-Marie-Tooth disease, type 2 | (6) |
| R162P | Motor | Spastic Paraplegia | (7) |
| R162W | Motor | Spastic Paraplegia | (8, 9) |
| S189P | Motor | Charcot-Marie-Tooth disease, type 2 | (1, 2) |
| A194P | Motor | Spastic Paraplegia | (10) |
| T196N | Motor | Charcot-Marie-Tooth disease, type 2 | (11) |
| M198T | Motor | Spastic Paraplegia | (3) |
| S202N | Motor (Sw1) | Spastic Paraplegia | (12) |
| S203C | Motor (Sw1) | Spastic Paraplegia | (13) |
| R204Q | Motor (Sw1) | Spastic paraplegia | (3, 14, 15) |
| R204P | Motor (Sw1) | Spastic paraplegia | (16) |
| R204W | Motor (Sw1) | Spastic paraplegia | (17-19) |
| V231L | Motor | Spastic paraplegia | (12) |
| D232N | Motor (Sw2) | Charcot-Marie-Tooth disease, type 2 | (20) |
| G235E | Motor (Sw2) | Spastic paraplegia | (12) |
| E237V | Motor (Sw2) | West syndrome and severe global developmental delay | (21) |
| L249V | Motor | Spastic paraplegia | (22) |
| E251K | Motor | Spastic paraplegia | (3) |
| K253N | Motor | Spastic paraplegia | (23, 24) |
| I255M | Motor | Spastic paraplegia | (25) |
| N256S | Motor | Spastic paraplegia | (23, 26, 27) |
| K257N | Motor | Spastic paraplegia | (3) |
| S258L | Motor | Spastic paraplegia | (28) |
| L259Q | Motor | Spastic paraplegia | (29) |
| L262P | Motor | Spastic paraplegia | (30) |
| A268T | Motor | Distal spinal muscular atrophy, adult-onset | (31, 32) |
| Y276C | Motor | Spastic paraplegia | (33) |
| P278L | Motor | Spastic paraplegia | (28) |
| R280C | Motor | Spastic paraplegia | (20, 23, 34) |
| R280H | Motor | Spastic paraplegia | (3) |
| R280L | Motor | Spastic paraplegia | (3) |
| T285I | Motor | Charcot-Marie-Tooth disease, type 2 | (6) |
| D290H | Motor | Spastic paraplegia | (35) |
| R297Q | Motor | Amyotrophic lateral sclerosis | (1) |
| R323W | Motor | Spastic paraplegia | (36, 37) |
| Q341R | Neck coil *c* | Spastic paraplegia | (38) |
| K362N | Neck coil *c* | Spastic paraplegia | (39) |
| A391V | Hinge 1 | Spastic paraplegia | (16) |
| E413G | Hinge 1 | Amyotrophic lateral sclerosis | (40) |
| K416N | CC1 *c* | Ataxic neuropathy | (41) |
| R423H | CC1 *c* | Amyotrophic lateral sclerosis | (42) |
| R468W | CC1 *f* | Spastic paraplegia | (43) |
| Q474H | CC1 *e* | Amyotrophic lateral sclerosis | (40) |
| L494M | CC1 *d* | Charcot-Marie-Tooth disease, type 2 | (42) |
| H542N | CC1 *c* | Amyotrophic lateral sclerosis | (1) |
| L558P | Hinge 2 | Charcot-Marie-Tooth disease, type 2 | (18) |
| G568R | Hinge 2 | Spastic paraplegia | (44) |
| S577G | Hinge 2 | Amyotrophic lateral sclerosis | (40) |
| A579T | Hinge 2 | Amyotrophic lateral sclerosis | (45) |

| | | | |
|---|---|---|---|
| T585A | Hinge 2 | Amyotrophic lateral sclerosis | (42) |
| R588Q | Hinge 2 | Amyotrophic lateral sclerosis | (42) |
| R606W | CC2 *b* | Spastic paraplegia | (46) |
| A669T | CC2 *b* | Charcot-Marie-Tooth disease, type 2 | (47) |
| R716Q | CC2 *g* | Amyotrophic lateral sclerosis | (42) |
| R716W | CC2 *g* | Spastic paraplegia | (48) |
| R718W | CC2 *b* | Amyotrophic lateral sclerosis | (4) |
| E755K | CC2 *d* | Mitochondrial disease | (49) |
| E758K | CC2 *a* | Spastic paraplegia | (12, 50, 51) |
| Q764* | CC2 *f* | Charcot-Marie-Tooth disease, type 2 | (52) |
| E785G | KLC binding | Peripheral neuropathy | (41) |
| E818Q | KLC binding | Ataxia | (53) |
| D853N | Cargo binding | Amyotrophic lateral sclerosis | (42) |
| E881K | Cargo binding | Amyotrophic lateral sclerosis | (42) |
| K907M | Auxillary MT binding site | Leber Optic Myopathy | (54) |
| V922I | IAK motif | Amyotrophic lateral sclerosis | (55) |
| T976I | C-terminal tail | Amyotrophic lateral sclerosis | (56) |
| A980V | C-terminal tail | Spastic paraplegia | (57) |
| P986L | C-terminal tail | Amyotrophic lateral sclerosis | (40, 58) |
| D1002G | C-terminal tail | Amyotrophic lateral sclerosis | (59) |
| R1007G | C-terminal tail | Amyotrophic lateral sclerosis | (58) |
| R1007K | C-terminal tail | Amyotrophic lateral sclerosis | (58) |
| F1023C | C-terminal tail | Amyotrophic lateral sclerosis | (55) |
| E1028D | C-terminal tail | Amyotrophic lateral sclerosis | (42) |

Note: in addition, 15 splicing mutations have been reported of which the majority (12) cause Amyotrophic lateral sclerosis of which most affect splicing of exon 27 (40) and results in misregulation of Kif5a, with oligomer formation, and abolishment of autoinhibition resulting in a toxic gain of function (60, 61) as skipping of exon 27 leads to a novel 39 amino acid sequence at the C-terminal region of Kif5a, following the conserved IAK motif.

**Supplemental Table 2: Crosslinking Mass Spectrometry data for WT KIF5A and for the exon 27 mutant KIF5A.** Crosslinks within the cut off distance of 27Å are shown.

WT — This table shows crosslinks observed for all 5 experiments

| Linked Res1 | Seqpos1 | Chain | Linked Res2 | Seqpos2 | Chain | Distance (Å) |
|---|---|---|---|---|---|---|
| LYS | 45 | A | LYS | 241 | A | 23.3 |
| LYS | 45 | B | LYS | 241 | B | 23.5 |
| LYS | 45 | A | SER | 240 | A | 24.6 |
| LYS | 45 | B | SER | 240 | B | 24.7 |
| TYR | 47 | A | LYS | 241 | A | 23.5 |
| TYR | 47 | B | LYS | 241 | B | 23.9 |
| THR | 93 | A | LYS | 188 | A | 12.4 |
| THR | 93 | B | LYS | 188 | B | 12.4 |
| THR | 95 | B | LYS | 188 | B | 16.4 |
| THR | 95 | A | LYS | 188 | A | 16.5 |
| LYS | 99 | A | LYS | 188 | A | 13 |
| LYS | 99 | B | LYS | 188 | B | 13.2 |
| LYS | 99 | A | SER | 189 | A | 15.1 |
| LYS | 99 | B | SER | 189 | B | 15.5 |
| LYS | 99 | B | LYS | 907 | B | 21.7 |
| LYS | 99 | B | LYS | 901 | B | 23.6 |
| LYS | 99 | A | LYS | 901 | B | 25.1 |
| LYS | 99 | A | LYS | 907 | B | 26.5 |
| TYR | 139 | A | LYS | 241 | A | 19.5 |
| TYR | 139 | B | LYS | 241 | B | 19.6 |
| LYS | 142 | A | LYS | 283 | A | 7.8 |
| LYS | 142 | B | LYS | 283 | B | 7.8 |
| LYS | 142 | A | THR | 196 | A | 17.7 |
| LYS | 142 | B | THR | 196 | B | 17.7 |
| LYS | 142 | A | THR | 242 | A | 21.9 |
| LYS | 142 | B | THR | 242 | B | 22 |
| LYS | 142 | A | LYS | 241 | A | 24 |
| LYS | 142 | B | LYS | 241 | B | 24 |
| THR | 150 | B | LYS | 283 | B | 18.7 |
| THR | 150 | A | LYS | 283 | A | 19.1 |
| THR | 150 | B | LYS | 253 | B | 26.2 |
| THR | 150 | A | LYS | 253 | A | 26.6 |
| LYS | 151 | A | THR | 170 | A | 8.8 |
| LYS | 151 | B | THR | 170 | B | 9 |
| THR | 152 | A | LYS | 426 | A | 12.8 |
| THR | 152 | A | LYS | 426 | B | 14.4 |
| SER | 155 | A | LYS | 283 | A | 7.8 |
| SER | 155 | B | LYS | 283 | B | 8.2 |
| SER | 155 | A | LYS | 416 | A | 11.9 |
| SER | 155 | A | LYS | 416 | B | 17.8 |
| LYS | 160 | A | LYS | 416 | B | 14.2 |
| LYS | 160 | A | LYS | 345 | B | 14.2 |
| LYS | 160 | A | THR | 220 | B | 15.7 |
| LYS | 160 | A | LYS | 416 | A | 16 |
| LYS | 160 | A | LYS | 223 | A | 17 |
| LYS | 160 | B | LYS | 223 | B | 17 |

Exon27 Mutant — This table shows crosslinks observed for all 3 experiments

| LinkedRes1 | Seqpos1 | Chain | Linked Res2 | Seqpos2 | Chain | Distance (Å) |
|---|---|---|---|---|---|---|
| LYS | 45 | A | SER | 240 | A | 24.6 |
| LYS | 45 | B | SER | 240 | B | 24.7 |
| LYS | 45 | A | THR | 242 | A | 26.7 |
| LYS | 45 | B | THR | 242 | B | 26.9 |
| TYR | 47 | A | LYS | 241 | A | 23.5 |
| TYR | 47 | B | LYS | 241 | B | 23.9 |
| THR | 93 | A | LYS | 188 | A | 12.4 |
| THR | 93 | B | LYS | 188 | B | 12.4 |
| THR | 95 | B | LYS | 188 | B | 16.4 |
| THR | 95 | A | LYS | 188 | A | 16.5 |
| LYS | 99 | A | LYS | 188 | A | 13 |
| LYS | 99 | B | LYS | 188 | B | 13.2 |
| LYS | 99 | A | SER | 189 | A | 15.1 |
| LYS | 99 | B | SER | 189 | B | 15.5 |
| LYS | 99 | A | SER | 202 | A | 16.2 |
| LYS | 99 | B | SER | 202 | B | 16.3 |
| LYS | 99 | A | SER | 203 | A | 17.7 |
| LYS | 99 | B | SER | 203 | B | 17.7 |
| LYS | 99 | B | LYS | 907 | B | 21.7 |
| LYS | 99 | A | THR | 196 | A | 22.5 |
| LYS | 99 | B | THR | 196 | B | 22.9 |
| LYS | 99 | B | LYS | 901 | B | 23.6 |
| LYS | 99 | A | LYS | 901 | B | 25.1 |
| LYS | 99 | A | LYS | 907 | B | 26.5 |
| TYR | 121 | B | LYS | 444 | B | 16.9 |
| TYR | 139 | A | LYS | 241 | A | 19.5 |
| TYR | 139 | B | LYS | 241 | B | 19.6 |
| LYS | 142 | A | LYS | 283 | A | 7.8 |
| LYS | 142 | B | LYS | 283 | B | 7.8 |
| LYS | 142 | A | LYS | 253 | A | 14.8 |
| LYS | 142 | B | LYS | 253 | B | 14.9 |
| LYS | 142 | A | SER | 260 | A | 15.6 |
| LYS | 142 | B | SER | 260 | B | 15.6 |
| LYS | 142 | A | TYR | 279 | A | 15.7 |
| LYS | 142 | B | TYR | 279 | B | 15.9 |
| LYS | 142 | A | THR | 196 | A | 17.7 |
| LYS | 142 | B | THR | 196 | B | 17.7 |
| LYS | 142 | A | THR | 242 | A | 21.9 |
| LYS | 142 | B | THR | 242 | B | 22 |
| LYS | 142 | A | LYS | 241 | A | 24 |
| LYS | 142 | B | LYS | 241 | B | 24 |
| THR | 150 | B | LYS | 283 | B | 18.7 |
| THR | 150 | A | LYS | 283 | A | 19.1 |
| THR | 150 | B | LYS | 253 | B | 26.2 |
| THR | 150 | A | LYS | 253 | A | 26.6 |
| THR | 152 | A | LYS | 426 | A | 6.7 |

| | | | | | | |
|---|---|---|---|---|---|---|
| LYS | 160 | A | LYS | 345 | A | 17.2 |
| LYS | 160 | B | LYS | 283 | B | 18 |
| LYS | 160 | A | LYS | 283 | A | 18.5 |
| LYS | 160 | B | THR | 285 | B | 18.7 |
| LYS | 160 | A | THR | 285 | A | 19.1 |
| LYS | 160 | A | LYS | 223 | B | 20 |
| LYS | 160 | B | LYS | 345 | B | 22 |
| LYS | 160 | A | THR | 220 | A | 24.6 |
| LYS | 160 | B | THR | 220 | B | 24.6 |
| LYS | 167 | A | SER | 155 | A | 5.8 |
| LYS | 167 | B | SER | 155 | B | 6.2 |
| LYS | 167 | A | LYS | 283 | A | 11.7 |
| LYS | 167 | B | LYS | 283 | B | 12 |
| LYS | 167 | B | LYS | 911 | B | 22.5 |
| LYS | 167 | B | LYS | 907 | B | 23.7 |
| LYS | 188 | B | LYS | 907 | B | 16.4 |
| THR | 196 | A | LYS | 241 | A | 22.4 |
| THR | 196 | B | LYS | 241 | B | 22.4 |
| SER | 202 | A | LYS | 241 | A | 17.1 |
| SER | 202 | B | LYS | 241 | B | 17.1 |
| LYS | 214 | B | LYS | 223 | B | 6.7 |
| LYS | 214 | A | LYS | 223 | A | 7 |
| LYS | 214 | A | LYS | 223 | B | 16.8 |
| LYS | 214 | B | LYS | 160 | B | 17.2 |
| LYS | 214 | A | LYS | 160 | A | 17.4 |
| LYS | 214 | A | LYS | 160 | B | 23.2 |
| LYS | 214 | A | SER | 155 | A | 23.6 |
| LYS | 214 | B | SER | 155 | B | 23.7 |
| LYS | 223 | A | SER | 155 | B | 26.7 |
| LYS | 227 | B | LYS | 283 | B | 17.7 |
| LYS | 227 | A | LYS | 283 | A | 17.8 |
| LYS | 227 | B | SER | 282 | B | 20.4 |
| LYS | 227 | A | SER | 282 | A | 20.5 |
| LYS | 238 | A | SER | 202 | A | 16 |
| LYS | 238 | B | SER | 202 | B | 16 |
| LYS | 238 | A | THR | 196 | A | 23.1 |
| LYS | 238 | B | THR | 196 | B | 23.2 |
| LYS | 253 | A | TYR | 139 | A | 12.4 |
| LYS | 253 | B | TYR | 139 | B | 12.4 |
| LYS | 253 | A | SER | 282 | A | 13.7 |
| LYS | 253 | B | SER | 282 | B | 13.7 |
| LYS | 253 | A | SER | 203 | A | 14 |
| LYS | 253 | B | SER | 203 | B | 14 |
| LYS | 253 | A | LYS | 283 | A | 16.2 |
| LYS | 253 | B | LYS | 283 | B | 16.2 |
| LYS | 253 | A | THR | 196 | A | 20.8 |
| LYS | 253 | B | THR | 196 | B | 20.8 |
| LYS | 257 | A | SER | 236 | A | 10.9 |
| LYS | 257 | B | SER | 236 | B | 10.9 |
| LYS | 257 | A | SER | 202 | A | 15.9 |
| LYS | 257 | B | SER | 202 | B | 15.9 |
| SER | 258 | A | LYS | 238 | A | 12.8 |
| SER | 258 | B | LYS | 238 | B | 12.8 |

| | | | | | | |
|---|---|---|---|---|---|---|
| THR | 152 | A | LYS | 167 | A | 11 |
| THR | 152 | B | LYS | 167 | B | 11.9 |
| THR | 152 | A | LYS | 283 | A | 13.3 |
| THR | 152 | B | LYS | 283 | B | 13.5 |
| THR | 152 | A | LYS | 426 | B | 15.1 |
| SER | 155 | A | LYS | 167 | A | 5.8 |
| SER | 155 | B | LYS | 167 | B | 6.2 |
| SER | 155 | A | LYS | 426 | A | 7.5 |
| SER | 155 | A | LYS | 283 | A | 7.8 |
| SER | 155 | B | LYS | 283 | B | 8.2 |
| SER | 155 | A | LYS | 431 | A | 11.4 |
| SER | 155 | A | LYS | 426 | B | 12.4 |
| SER | 155 | A | LYS | 431 | B | 14.2 |
| LYS | 160 | A | LYS | 416 | B | 14.2 |
| LYS | 160 | A | LYS | 426 | B | 14.5 |
| LYS | 160 | A | THR | 220 | B | 15.7 |
| LYS | 160 | A | LYS | 416 | A | 16 |
| LYS | 160 | A | LYS | 223 | A | 17 |
| LYS | 160 | B | LYS | 223 | B | 17 |
| LYS | 160 | B | LYS | 214 | B | 17.2 |
| LYS | 160 | A | LYS | 214 | A | 17.4 |
| LYS | 160 | B | LYS | 283 | B | 18 |
| LYS | 160 | A | LYS | 283 | A | 18.5 |
| LYS | 160 | A | LYS | 223 | B | 20 |
| LYS | 160 | A | LYS | 426 | A | 22.4 |
| LYS | 160 | A | THR | 220 | A | 24.6 |
| LYS | 160 | B | THR | 220 | B | 24.6 |
| LYS | 160 | A | LYS | 214 | B | 24.8 |
| LYS | 167 | A | LYS | 426 | B | 7.3 |
| LYS | 167 | A | LYS | 426 | A | 10.7 |
| LYS | 167 | A | LYS | 283 | A | 11.7 |
| LYS | 167 | B | LYS | 283 | B | 12 |
| LYS | 167 | B | TYR | 927 | B | 13.6 |
| LYS | 167 | A | LYS | 416 | A | 15.3 |
| LYS | 167 | A | THR | 220 | B | 15.6 |
| LYS | 167 | A | LYS | 416 | B | 16.8 |
| LYS | 167 | B | LYS | 911 | B | 22.5 |
| LYS | 167 | A | LYS | 920 | B | 23.5 |
| LYS | 167 | B | LYS | 907 | B | 23.7 |
| THR | 170 | A | LYS | 426 | B | 9.5 |
| THR | 170 | A | LYS | 426 | A | 15.2 |
| LYS | 188 | A | SER | 207 | A | 10 |
| LYS | 188 | B | SER | 207 | B | 10 |
| LYS | 188 | B | TYR | 893 | B | 24.4 |
| THR | 196 | A | LYS | 253 | A | 20.8 |
| THR | 196 | B | LYS | 253 | B | 20.8 |
| THR | 196 | A | LYS | 241 | A | 22.4 |
| THR | 196 | B | LYS | 241 | B | 22.4 |
| THR | 196 | A | LYS | 257 | A | 22.5 |
| THR | 196 | B | LYS | 257 | B | 22.5 |
| SER | 202 | A | LYS | 257 | A | 15.9 |
| SER | 202 | B | LYS | 257 | B | 15.9 |
| SER | 202 | A | LYS | 238 | A | 16 |

| | | | | | | |
|---|---|---|---|---|---|---|
| TYR | 279 | A | LYS | 416 | A | 14.1 |
| TYR | 279 | A | LYS | 416 | B | 25.4 |
| SER | 305 | A | LYS | 238 | A | 15 |
| SER | 305 | B | LYS | 238 | B | 15 |
| SER | 307 | A | LYS | 241 | A | 15.2 |
| SER | 307 | B | LYS | 241 | B | 15.2 |
| SER | 307 | B | LYS | 238 | B | 16.8 |
| SER | 307 | A | LYS | 238 | A | 16.9 |
| THR | 314 | A | LYS | 241 | A | 13.2 |
| THR | 314 | B | LYS | 241 | B | 13.2 |
| LYS | 315 | A | SER | 236 | A | 11.7 |
| LYS | 315 | B | SER | 236 | B | 11.7 |
| LYS | 315 | A | THR | 242 | A | 17.7 |
| LYS | 315 | B | THR | 242 | B | 17.7 |
| LYS | 352 | A | LYS | 350 | A | 5.4 |
| LYS | 352 | B | LYS | 350 | B | 5.5 |
| LYS | 352 | A | LYS | 350 | B | 9.2 |
| THR | 353 | A | LYS | 352 | A | 3.8 |
| THR | 353 | B | LYS | 352 | B | 3.8 |
| THR | 353 | A | LYS | 352 | B | 7.6 |
| LYS | 354 | A | THR | 353 | A | 3.8 |
| LYS | 354 | B | THR | 353 | B | 3.8 |
| LYS | 354 | A | THR | 353 | B | 8.8 |
| LYS | 416 | A | LYS | 283 | A | 13.7 |
| LYS | 416 | A | LYS | 167 | A | 15.3 |
| LYS | 416 | A | THR | 285 | A | 16.7 |
| LYS | 416 | B | LYS | 357 | B | 23.7 |
| TYR | 417 | A | LYS | 283 | A | 14.9 |
| LYS | 426 | A | THR | 152 | A | 6.7 |
| LYS | 426 | A | SER | 155 | A | 7.5 |
| LYS | 426 | A | LYS | 167 | A | 10.7 |
| LYS | 426 | A | LYS | 283 | A | 13.6 |
| LYS | 426 | A | THR | 170 | A | 15.2 |
| LYS | 426 | A | SER | 282 | A | 15.9 |
| LYS | 426 | A | LYS | 160 | A | 22.4 |
| LYS | 431 | A | SER | 258 | A | 21.9 |
| LYS | 431 | B | LYS | 446 | B | 22.3 |
| LYS | 431 | A | LYS | 446 | A | 23 |
| LYS | 431 | A | LYS | 446 | B | 23.1 |
| SER | 439 | B | LYS | 446 | B | 10.5 |
| SER | 439 | A | LYS | 446 | A | 10.6 |
| SER | 439 | A | LYS | 446 | B | 13.9 |
| LYS | 444 | B | LYS | 901 | B | 19.7 |
| LYS | 444 | B | LYS | 907 | B | 20 |
| LYS | 444 | A | THR | 459 | A | 22.6 |
| LYS | 444 | B | THR | 459 | B | 22.6 |
| LYS | 444 | A | THR | 459 | B | 24.3 |
| LYS | 446 | B | LYS | 907 | B | 24.8 |
| LYS | 465 | B | LYS | 888 | B | 20 |
| LYS | 465 | A | LYS | 888 | B | 22.9 |
| LYS | 465 | B | LYS | 873 | B | 23.2 |
| LYS | 465 | A | LYS | 873 | B | 25.3 |
| LYS | 508 | A | LYS | 845 | A | 12.4 |

| | | | | | | |
|---|---|---|---|---|---|---|
| SER | 202 | B | LYS | 238 | B | 16 |
| SER | 202 | A | LYS | 241 | A | 17.1 |
| SER | 202 | B | LYS | 241 | B | 17.1 |
| SER | 203 | A | LYS | 241 | A | 13.5 |
| SER | 203 | B | LYS | 241 | B | 13.5 |
| LYS | 214 | B | THR | 220 | B | 14.4 |
| LYS | 214 | A | THR | 220 | A | 14.8 |
| LYS | 214 | A | THR | 220 | B | 19.5 |
| THR | 220 | A | THR | 220 | B | 26.6 |
| LYS | 227 | B | LYS | 283 | B | 17.7 |
| LYS | 227 | A | LYS | 283 | A | 17.8 |
| TYR | 229 | A | LYS | 315 | A | 26.4 |
| TYR | 229 | B | LYS | 315 | B | 26.4 |
| SER | 236 | A | LYS | 315 | A | 11.7 |
| SER | 236 | B | LYS | 315 | B | 11.7 |
| SER | 236 | A | LYS | 283 | A | 17.1 |
| SER | 236 | B | LYS | 283 | B | 17.2 |
| LYS | 238 | A | SER | 258 | A | 12.8 |
| LYS | 238 | B | SER | 258 | B | 12.8 |
| LYS | 238 | A | SER | 305 | A | 15 |
| LYS | 238 | B | SER | 305 | B | 15 |
| LYS | 238 | B | SER | 307 | B | 16.8 |
| LYS | 238 | A | SER | 307 | A | 16.9 |
| SER | 240 | A | LYS | 257 | A | 14.5 |
| SER | 240 | B | LYS | 257 | B | 14.5 |
| LYS | 241 | A | THR | 314 | A | 13.2 |
| LYS | 241 | B | THR | 314 | B | 13.2 |
| LYS | 241 | A | LYS | 283 | A | 25.3 |
| LYS | 241 | B | LYS | 283 | B | 25.3 |
| THR | 242 | A | LYS | 315 | A | 17.7 |
| THR | 242 | B | LYS | 315 | B | 17.7 |
| LYS | 253 | A | SER | 282 | A | 13.7 |
| LYS | 253 | B | SER | 282 | B | 13.7 |
| LYS | 253 | A | LYS | 283 | A | 16.2 |
| LYS | 253 | B | LYS | 283 | B | 16.2 |
| LYS | 253 | A | TYR | 279 | A | 19 |
| LYS | 253 | B | TYR | 279 | B | 19 |
| LYS | 253 | A | LYS | 426 | A | 22.3 |
| LYS | 253 | A | SER | 305 | A | 23.1 |
| LYS | 253 | B | SER | 305 | B | 23.1 |
| LYS | 253 | A | LYS | 431 | A | 26.3 |
| LYS | 257 | A | LYS | 283 | A | 10.3 |
| LYS | 257 | B | LYS | 283 | B | 10.3 |
| SER | 258 | A | LYS | 431 | A | 21.9 |
| SER | 258 | A | LYS | 431 | B | 25.9 |
| THR | 273 | A | LYS | 416 | A | 23.8 |
| LYS | 274 | B | SER | 332 | B | 18 |
| LYS | 274 | A | SER | 332 | A | 18.2 |
| TYR | 279 | A | LYS | 416 | A | 14.1 |
| TYR | 279 | A | LYS | 416 | B | 25.4 |
| SER | 282 | A | LYS | 416 | A | 14 |
| SER | 282 | A | LYS | 416 | B | 25.7 |
| LYS | 283 | A | LYS | 426 | A | 13.6 |

| | | | | | | |
|---|---|---|---|---|---|---|
| LYS | 508 | A | LYS | 845 | B | 19.8 |
| LYS | 508 | B | LYS | 845 | B | 24.5 |
| LYS | 522 | A | SER | 820 | A | 17.1 |
| LYS | 522 | A | SER | 820 | B | 23.1 |
| LYS | 522 | B | SER | 820 | B | 26.2 |
| LYS | 545 | B | LYS | 799 | B | 16.6 |
| LYS | 545 | A | LYS | 799 | A | 20.5 |
| LYS | 556 | B | THR | 806 | B | 23.3 |
| LYS | 593 | B | LYS | 777 | B | 17.8 |
| LYS | 593 | A | LYS | 799 | A | 20 |
| LYS | 593 | A | LYS | 777 | A | 22.4 |
| LYS | 593 | B | LYS | 799 | B | 24.9 |
| LYS | 593 | A | LYS | 777 | B | 26.6 |
| SER | 634 | A | LYS | 732 | B | 23.4 |
| SER | 634 | A | LYS | 732 | A | 26 |
| SER | 634 | B | LYS | 732 | B | 27 |
| LYS | 639 | A | LYS | 732 | B | 15.6 |
| LYS | 639 | A | LYS | 732 | A | 20.1 |
| LYS | 639 | B | LYS | 732 | B | 20.9 |
| LYS | 639 | A | THR | 727 | A | 22.5 |
| LYS | 639 | A | THR | 727 | B | 24 |
| TYR | 646 | A | LYS | 732 | B | 9.3 |
| TYR | 646 | A | LYS | 732 | A | 15.8 |
| TYR | 646 | B | LYS | 732 | B | 16 |
| SER | 649 | A | LYS | 653 | A | 6.6 |
| SER | 649 | B | LYS | 653 | B | 6.6 |
| SER | 649 | A | LYS | 732 | B | 11.2 |
| SER | 649 | A | LYS | 653 | B | 14.2 |
| SER | 649 | B | LYS | 732 | B | 15.2 |
| SER | 649 | A | LYS | 732 | A | 17.2 |
| LYS | 653 | A | LYS | 724 | A | 13.2 |
| LYS | 653 | A | LYS | 732 | B | 13.3 |
| LYS | 653 | B | LYS | 732 | B | 16.1 |
| LYS | 653 | A | LYS | 724 | B | 16.1 |
| LYS | 653 | A | LYS | 732 | A | 19.4 |
| LYS | 653 | B | LYS | 724 | B | 19.6 |
| TYR | 661 | A | LYS | 653 | A | 12.2 |
| TYR | 661 | B | LYS | 653 | B | 12.2 |
| TYR | 661 | A | LYS | 653 | B | 13.4 |
| LYS | 696 | A | LYS | 697 | A | 3.9 |
| LYS | 696 | B | LYS | 697 | B | 3.9 |
| LYS | 696 | A | LYS | 697 | B | 13.9 |
| THR | 727 | A | LYS | 732 | A | 8.5 |
| THR | 727 | B | LYS | 732 | B | 8.5 |
| THR | 727 | A | LYS | 732 | B | 10 |
| LYS | 737 | A | LYS | 639 | A | 16.2 |
| LYS | 737 | B | LYS | 639 | B | 16.7 |
| LYS | 737 | A | LYS | 639 | B | 25.7 |
| TYR | 749 | A | LYS | 618 | A | 23.6 |
| TYR | 749 | B | LYS | 618 | B | 24.6 |
| LYS | 751 | B | THR | 622 | B | 15 |
| LYS | 751 | A | THR | 622 | A | 21 |
| LYS | 751 | A | THR | 622 | B | 26 |

| | | | | | | |
|---|---|---|---|---|---|---|
| LYS | 283 | A | LYS | 416 | A | 13.7 |
| LYS | 283 | A | LYS | 426 | B | 19 |
| LYS | 283 | A | LYS | 416 | B | 23.6 |
| LYS | 350 | A | LYS | 352 | A | 5.4 |
| LYS | 350 | B | LYS | 352 | B | 5.5 |
| LYS | 350 | B | LYS | 354 | B | 6.5 |
| LYS | 350 | A | LYS | 354 | A | 6.6 |
| LYS | 350 | A | LYS | 352 | B | 9.7 |
| LYS | 350 | A | LYS | 354 | B | 12 |
| LYS | 352 | A | THR | 353 | A | 3.8 |
| LYS | 352 | B | THR | 353 | B | 3.8 |
| LYS | 352 | A | THR | 353 | B | 7.3 |
| LYS | 357 | B | LYS | 416 | B | 23.7 |
| LYS | 357 | A | LYS | 416 | B | 25.6 |
| LYS | 416 | A | LYS | 416 | B | 14.1 |
| LYS | 431 | B | LYS | 446 | B | 22.3 |
| LYS | 431 | A | LYS | 446 | A | 23 |
| LYS | 431 | A | LYS | 446 | B | 23.1 |
| SER | 439 | B | LYS | 446 | B | 10.5 |
| SER | 439 | A | LYS | 446 | A | 10.6 |
| SER | 439 | A | LYS | 446 | B | 13.9 |
| LYS | 444 | B | LYS | 901 | B | 19.7 |
| LYS | 444 | B | LYS | 907 | B | 20 |
| LYS | 446 | B | LYS | 901 | B | 23.1 |
| LYS | 446 | B | LYS | 907 | B | 24.8 |
| LYS | 465 | A | LYS | 465 | B | 13.8 |
| LYS | 465 | B | LYS | 883 | B | 19.4 |
| LYS | 465 | B | LYS | 888 | B | 20 |
| LYS | 465 | A | LYS | 888 | B | 22.9 |
| LYS | 465 | B | LYS | 873 | B | 23.2 |
| LYS | 465 | A | LYS | 883 | B | 24.6 |
| LYS | 465 | A | LYS | 873 | B | 25.3 |
| LYS | 465 | B | TYR | 893 | B | 26.9 |
| LYS | 508 | A | THR | 841 | A | 9.8 |
| LYS | 508 | A | LYS | 845 | A | 12.4 |
| LYS | 508 | A | THR | 841 | B | 17.1 |
| LYS | 508 | A | LYS | 845 | B | 19.8 |
| LYS | 508 | B | THR | 841 | B | 22.3 |
| LYS | 508 | B | LYS | 845 | B | 24.5 |
| SER | 520 | A | LYS | 842 | A | 23 |
| LYS | 522 | A | SER | 820 | A | 17.1 |
| LYS | 522 | A | SER | 820 | B | 23.1 |
| LYS | 522 | B | SER | 820 | B | 26.2 |
| SER | 540 | A | LYS | 799 | A | 20 |
| SER | 540 | B | LYS | 799 | B | 22.7 |
| SER | 540 | A | LYS | 799 | B | 25.1 |
| LYS | 545 | B | LYS | 799 | B | 16.6 |
| LYS | 545 | A | LYS | 799 | A | 20.5 |
| LYS | 556 | B | THR | 806 | B | 23.3 |
| LYS | 556 | B | THR | 807 | B | 26.6 |
| LYS | 593 | B | THR | 786 | B | 12.1 |
| LYS | 593 | A | THR | 786 | A | 15.1 |
| LYS | 593 | B | LYS | 781 | B | 15.4 |

| | | | | | | |
|---|---|---|---|---|---|---|
| LYS | 753 | B | THR | 622 | B | 15.9 |
| LYS | 753 | A | THR | 622 | A | 16.7 |
| LYS | 753 | A | LYS | 618 | A | 19.5 |
| LYS | 753 | B | LYS | 618 | B | 19.9 |
| LYS | 753 | A | THR | 622 | B | 22.4 |
| LYS | 753 | A | LYS | 618 | B | 26.1 |
| SER | 754 | B | LYS | 618 | B | 16.5 |
| SER | 754 | A | LYS | 618 | A | 20.7 |
| LYS | 759 | B | SER | 754 | B | 8.4 |
| LYS | 759 | A | SER | 754 | A | 8.7 |
| LYS | 759 | A | SER | 754 | B | 12.8 |
| LYS | 759 | A | LYS | 618 | A | 14.2 |
| LYS | 759 | B | LYS | 618 | B | 15.5 |
| LYS | 759 | A | LYS | 618 | B | 22 |
| SER | 760 | A | LYS | 753 | A | 10.5 |
| SER | 760 | B | LYS | 753 | B | 10.5 |
| SER | 760 | A | LYS | 753 | B | 13.4 |
| LYS | 762 | B | LYS | 618 | B | 12.9 |
| LYS | 762 | B | THR | 622 | B | 13.8 |
| LYS | 762 | A | LYS | 618 | A | 14.8 |
| LYS | 762 | A | THR | 622 | A | 16.9 |
| LYS | 762 | A | THR | 622 | B | 23 |
| LYS | 762 | A | LYS | 618 | B | 23.4 |
| THR | 767 | A | LYS | 777 | A | 15.2 |
| THR | 767 | B | LYS | 777 | B | 15.2 |
| THR | 767 | A | LYS | 777 | B | 17.8 |
| SER | 776 | B | LYS | 593 | B | 16.8 |
| SER | 776 | A | LYS | 593 | B | 24.4 |
| SER | 776 | A | LYS | 593 | A | 25.2 |
| LYS | 777 | A | LYS | 603 | A | 9.8 |
| LYS | 777 | B | LYS | 603 | B | 17.3 |
| LYS | 777 | A | LYS | 603 | B | 18.5 |
| LYS | 781 | B | LYS | 777 | B | 6.2 |
| LYS | 781 | A | LYS | 777 | A | 6.3 |
| LYS | 781 | A | SER | 776 | A | 8.6 |
| LYS | 781 | B | SER | 776 | B | 8.6 |
| LYS | 781 | A | LYS | 777 | B | 12.8 |
| LYS | 781 | A | SER | 776 | B | 14.5 |
| LYS | 781 | B | LYS | 593 | B | 15.4 |
| LYS | 781 | A | LYS | 593 | A | 17 |
| LYS | 781 | A | LYS | 593 | B | 18 |
| LYS | 781 | A | LYS | 799 | A | 26.1 |
| LYS | 781 | B | LYS | 799 | B | 26.8 |
| THR | 786 | B | LYS | 593 | B | 12.1 |
| THR | 786 | A | LYS | 593 | A | 15.1 |
| THR | 786 | A | LYS | 593 | B | 20.3 |
| LYS | 799 | A | SER | 540 | B | 16.2 |
| LYS | 799 | A | SER | 592 | A | 16.5 |
| LYS | 799 | A | SER | 540 | A | 20 |
| LYS | 799 | A | SER | 592 | B | 22.6 |
| LYS | 799 | B | SER | 540 | B | 22.7 |
| LYS | 799 | A | LYS | 599 | A | 24.9 |
| LYS | 799 | B | SER | 592 | B | 25.5 |

| | | | | | | |
|---|---|---|---|---|---|---|
| LYS | 593 | A | LYS | 781 | A | 17 |
| LYS | 593 | B | LYS | 777 | B | 17.8 |
| LYS | 593 | A | LYS | 593 | B | 17.9 |
| LYS | 593 | A | LYS | 799 | A | 20 |
| LYS | 593 | A | LYS | 777 | A | 22.4 |
| LYS | 593 | B | LYS | 799 | B | 24.9 |
| LYS | 593 | A | LYS | 781 | B | 26.2 |
| LYS | 593 | A | THR | 786 | B | 26.3 |
| LYS | 593 | A | LYS | 777 | B | 26.6 |
| LYS | 595 | A | LYS | 799 | A | 19.8 |
| SER | 596 | B | LYS | 777 | B | 17.2 |
| SER | 596 | A | LYS | 777 | A | 17.5 |
| SER | 596 | A | LYS | 777 | B | 22.6 |
| LYS | 599 | A | LYS | 603 | A | 6.1 |
| LYS | 599 | B | LYS | 603 | B | 6.1 |
| LYS | 599 | A | LYS | 777 | A | 12.8 |
| LYS | 599 | A | LYS | 603 | B | 14.7 |
| LYS | 599 | B | SER | 776 | B | 15.2 |
| LYS | 599 | A | SER | 776 | A | 15.9 |
| LYS | 599 | B | LYS | 777 | B | 17.5 |
| LYS | 599 | A | LYS | 777 | B | 18 |
| LYS | 599 | A | SER | 776 | B | 18.1 |
| LYS | 599 | A | LYS | 799 | A | 24.9 |
| SER | 600 | A | LYS | 777 | A | 14.2 |
| SER | 600 | B | LYS | 777 | B | 14.3 |
| SER | 600 | A | LYS | 777 | B | 19.8 |
| LYS | 603 | A | LYS | 777 | A | 9.8 |
| LYS | 603 | A | LYS | 603 | B | 13.8 |
| LYS | 603 | A | LYS | 777 | B | 16.2 |
| LYS | 603 | B | LYS | 777 | B | 17.3 |
| LYS | 603 | A | LYS | 618 | A | 22.7 |
| LYS | 603 | B | LYS | 618 | B | 22.8 |
| LYS | 603 | A | LYS | 618 | B | 25.4 |
| LYS | 618 | A | LYS | 759 | B | 9.3 |
| LYS | 618 | A | THR | 761 | B | 10.4 |
| LYS | 618 | A | LYS | 618 | B | 10.8 |
| LYS | 618 | A | THR | 767 | A | 11.2 |
| LYS | 618 | B | THR | 761 | B | 13.1 |
| LYS | 618 | A | SER | 754 | B | 14 |
| LYS | 618 | A | LYS | 759 | A | 14.2 |
| LYS | 618 | A | TYR | 770 | A | 14.6 |
| LYS | 618 | A | THR | 767 | B | 14.6 |
| LYS | 618 | A | THR | 761 | A | 14.9 |
| LYS | 618 | B | LYS | 759 | B | 15.5 |
| LYS | 618 | B | SER | 754 | B | 16.5 |
| LYS | 618 | A | TYR | 770 | B | 16.6 |
| LYS | 618 | A | LYS | 753 | B | 16.9 |
| LYS | 618 | A | LYS | 751 | B | 17.1 |
| LYS | 618 | A | LYS | 753 | A | 19.5 |
| LYS | 618 | B | THR | 767 | B | 19.5 |
| LYS | 618 | B | LYS | 753 | B | 19.9 |
| LYS | 618 | B | LYS | 751 | B | 20.2 |
| LYS | 618 | A | SER | 754 | A | 20.7 |

| | | | | | | |
|---|---|---|---|---|---|---|
| LYS | 811 | A | SER | 820 | B | 9.2 |
| LYS | 811 | B | SER | 820 | B | 13.2 |
| LYS | 811 | A | SER | 820 | A | 15.9 |
| LYS | 811 | A | SER | 825 | B | 18.6 |
| LYS | 811 | B | SER | 825 | B | 19.5 |
| LYS | 811 | A | SER | 825 | A | 20.8 |
| SER | 812 | B | LYS | 799 | B | 19.7 |
| SER | 812 | A | LYS | 799 | B | 21.8 |
| SER | 812 | A | LYS | 799 | A | 23.9 |
| SER | 820 | A | LYS | 829 | A | 16.4 |
| SER | 820 | B | LYS | 829 | B | 18.1 |
| SER | 820 | A | LYS | 829 | B | 21.6 |
| SER | 825 | A | LYS | 829 | A | 7.7 |
| SER | 825 | B | LYS | 829 | B | 8.2 |
| SER | 825 | A | LYS | 829 | B | 14.6 |
| LYS | 827 | A | SER | 825 | A | 5.6 |
| LYS | 827 | B | SER | 825 | B | 5.6 |
| LYS | 827 | A | SER | 825 | B | 13.2 |
| LYS | 827 | A | SER | 820 | A | 13.4 |
| LYS | 827 | B | SER | 820 | B | 13.5 |
| LYS | 827 | A | SER | 528 | B | 13.7 |
| LYS | 827 | A | SER | 528 | A | 16.7 |
| LYS | 827 | A | SER | 820 | B | 18 |
| LYS | 827 | B | THR | 841 | B | 20.8 |
| LYS | 827 | B | SER | 528 | B | 20.9 |
| LYS | 827 | A | THR | 841 | A | 21.3 |
| LYS | 827 | A | THR | 841 | B | 21.7 |
| LYS | 827 | B | LYS | 845 | B | 26.8 |
| THR | 841 | A | LYS | 845 | A | 6.1 |
| THR | 841 | B | LYS | 845 | B | 6.1 |
| THR | 841 | A | LYS | 845 | B | 12.4 |
| LYS | 842 | B | LYS | 845 | B | 5.2 |
| LYS | 842 | A | LYS | 845 | A | 5.4 |
| LYS | 842 | A | LYS | 845 | B | 13.5 |
| LYS | 860 | A | LYS | 873 | A | 20.3 |
| LYS | 860 | B | LYS | 873 | B | 20.3 |
| LYS | 860 | A | LYS | 873 | B | 21.4 |
| LYS | 883 | B | LYS | 888 | B | 8.7 |
| LYS | 883 | A | LYS | 888 | A | 8.8 |
| LYS | 883 | A | LYS | 888 | B | 13.7 |
| LYS | 883 | B | LYS | 465 | B | 19.4 |
| LYS | 911 | A | LYS | 907 | A | 12.1 |
| LYS | 911 | B | LYS | 907 | B | 14.4 |
| LYS | 911 | A | LYS | 907 | B | 15.7 |
| LYS | 920 | B | LYS | 907 | B | 11.5 |

| | | | | | | |
|---|---|---|---|---|---|---|
| LYS | 618 | A | TYR | 749 | B | 21.3 |
| LYS | 618 | B | TYR | 770 | B | 21.5 |
| LYS | 618 | A | LYS | 777 | A | 22.7 |
| LYS | 618 | A | TYR | 749 | A | 23.6 |
| LYS | 618 | A | LYS | 751 | A | 23.9 |
| LYS | 618 | B | TYR | 749 | B | 24.6 |
| LYS | 618 | A | LYS | 777 | B | 25.5 |
| THR | 622 | A | LYS | 762 | B | 9.9 |
| THR | 622 | A | LYS | 751 | B | 11.7 |
| THR | 622 | A | LYS | 753 | B | 12.4 |
| THR | 622 | B | LYS | 762 | B | 13.8 |
| THR | 622 | B | LYS | 751 | B | 15 |
| THR | 622 | B | LYS | 753 | B | 15.9 |
| THR | 622 | A | LYS | 753 | A | 16.7 |
| THR | 622 | A | LYS | 762 | A | 16.9 |
| THR | 622 | A | LYS | 751 | A | 21 |
| SER | 634 | A | LYS | 744 | B | 9.4 |
| SER | 634 | A | SER | 634 | B | 12.9 |
| SER | 634 | B | LYS | 744 | B | 13.2 |
| SER | 634 | A | LYS | 744 | A | 17.1 |
| SER | 634 | A | LYS | 737 | B | 17.9 |
| SER | 634 | A | LYS | 737 | A | 19.7 |
| SER | 634 | B | LYS | 737 | B | 20.5 |
| SER | 634 | A | LYS | 732 | B | 23.4 |
| SER | 634 | A | LYS | 732 | A | 26 |
| SER | 634 | B | LYS | 732 | B | 27 |
| LYS | 639 | A | LYS | 639 | B | 9.9 |
| LYS | 639 | A | LYS | 737 | B | 10.9 |
| LYS | 639 | A | LYS | 732 | B | 15.6 |
| LYS | 639 | A | LYS | 737 | A | 16.2 |
| LYS | 639 | B | LYS | 737 | B | 16.7 |
| LYS | 639 | A | LYS | 732 | A | 20.1 |
| LYS | 639 | B | LYS | 732 | B | 20.9 |
| LYS | 639 | A | THR | 727 | A | 22.5 |
| LYS | 639 | A | THR | 727 | B | 24 |
| SER | 649 | A | LYS | 732 | B | 11.2 |
| SER | 649 | B | LYS | 732 | B | 15.2 |
| SER | 649 | A | LYS | 732 | A | 17.2 |
| LYS | 653 | A | SER | 660 | A | 10.5 |
| LYS | 653 | B | SER | 660 | B | 10.5 |
| LYS | 653 | A | LYS | 653 | B | 11.2 |
| LYS | 653 | A | TYR | 661 | A | 12.2 |
| LYS | 653 | B | TYR | 661 | B | 12.2 |
| LYS | 653 | A | LYS | 724 | A | 13.2 |
| LYS | 653 | A | LYS | 732 | B | 13.3 |
| LYS | 653 | A | TYR | 661 | B | 13.3 |
| LYS | 653 | A | SER | 660 | B | 14.9 |
| LYS | 653 | A | SER | 663 | A | 15.4 |
| LYS | 653 | B | SER | 663 | B | 15.4 |
| LYS | 653 | A | LYS | 724 | B | 16.1 |
| LYS | 653 | B | LYS | 732 | B | 16.1 |
| LYS | 653 | A | SER | 663 | B | 18.5 |
| LYS | 653 | A | LYS | 732 | A | 19.4 |

| | | | | | | |
|---|---|---|---|---|---|---|
| LYS | 653 | B | LYS | 724 | B | 19.6 |
| LYS | 683 | B | LYS | 696 | B | 10.4 |
| LYS | 683 | A | THR | 689 | A | 12 |
| LYS | 683 | B | THR | 689 | B | 14.4 |
| LYS | 683 | B | SER | 705 | B | 18.8 |
| LYS | 683 | A | LYS | 696 | A | 19.4 |
| LYS | 683 | A | THR | 689 | B | 21.2 |
| LYS | 683 | A | LYS | 696 | B | 21.2 |
| LYS | 683 | A | SER | 705 | A | 22.3 |
| LYS | 696 | A | LYS | 697 | A | 3.9 |
| LYS | 696 | A | LYS | 697 | B | 13.9 |
| LYS | 732 | A | LYS | 732 | B | 9.2 |
| LYS | 753 | A | SER | 760 | A | 10.5 |
| LYS | 753 | B | SER | 760 | B | 10.5 |
| LYS | 753 | A | SER | 760 | B | 13.4 |
| TYR | 770 | A | LYS | 777 | A | 10.4 |
| TYR | 770 | B | LYS | 777 | B | 10.4 |
| TYR | 770 | A | LYS | 777 | B | 13 |
| TYR | 770 | B | LYS | 781 | B | 16.4 |
| TYR | 770 | A | LYS | 781 | A | 16.5 |
| TYR | 770 | A | LYS | 781 | B | 18.6 |
| SER | 776 | A | LYS | 777 | A | 3.8 |
| SER | 776 | B | LYS | 777 | B | 3.8 |
| SER | 776 | A | LYS | 781 | A | 8.6 |
| SER | 776 | B | LYS | 781 | B | 8.6 |
| SER | 776 | A | LYS | 777 | B | 11.5 |
| SER | 776 | A | LYS | 781 | B | 14.3 |
| LYS | 777 | B | LYS | 781 | B | 6.2 |
| LYS | 777 | A | LYS | 781 | A | 6.3 |
| LYS | 777 | A | LYS | 777 | B | 9.5 |
| LYS | 777 | A | LYS | 781 | B | 12.7 |
| LYS | 781 | A | GLN | 775 | A | 10 |
| LYS | 781 | B | GLN | 775 | B | 10 |
| LYS | 781 | A | GLN | 775 | B | 17.7 |
| LYS | 781 | A | LYS | 799 | A | 26.1 |
| LYS | 781 | B | LYS | 799 | B | 26.8 |
| LYS | 799 | A | LYS | 799 | B | 16.1 |
| LYS | 799 | B | SER | 812 | B | 19.7 |
| LYS | 799 | A | SER | 812 | A | 23.9 |
| LYS | 799 | A | SER | 812 | B | 25.9 |
| THR | 807 | B | LYS | 811 | B | 5.4 |
| THR | 807 | A | LYS | 811 | A | 11 |
| THR | 807 | A | LYS | 811 | B | 20.2 |
| LYS | 811 | A | SER | 812 | A | 3.8 |
| LYS | 811 | B | SER | 812 | B | 3.9 |
| LYS | 811 | A | SER | 812 | B | 8.4 |
| LYS | 811 | A | SER | 820 | B | 9.2 |
| LYS | 811 | B | SER | 820 | B | 13.2 |
| LYS | 811 | A | SER | 820 | A | 15.9 |
| SER | 820 | A | LYS | 829 | A | 16.4 |
| SER | 820 | B | LYS | 829 | B | 18.1 |
| SER | 820 | A | LYS | 829 | B | 21.6 |
| SER | 825 | A | LYS | 827 | A | 5.6 |

| | | | | | | |
|---|---|---|---|---|---|---|
| SER | 825 | B | LYS | 827 | B | 5.6 |
| SER | 825 | A | LYS | 829 | A | 7.7 |
| SER | 825 | B | LYS | 829 | B | 8.2 |
| SER | 825 | A | LYS | 827 | B | 11.8 |
| SER | 825 | A | LYS | 829 | B | 14.6 |
| LYS | 827 | B | THR | 841 | B | 20.8 |
| LYS | 827 | A | THR | 841 | A | 21.3 |
| LYS | 827 | A | THR | 841 | B | 21.7 |
| LYS | 827 | B | LYS | 845 | B | 26.8 |
| LYS | 842 | B | LYS | 845 | B | 5.2 |
| LYS | 842 | A | LYS | 845 | A | 5.4 |
| LYS | 842 | A | LYS | 845 | B | 13.5 |
| LYS | 845 | A | LYS | 845 | B | 12.8 |
| LYS | 845 | A | LYS | 860 | A | 23.4 |
| LYS | 845 | B | LYS | 860 | B | 23.4 |
| LYS | 845 | A | LYS | 860 | B | 26.2 |
| LYS | 860 | A | LYS | 873 | A | 20.3 |
| LYS | 860 | B | LYS | 873 | B | 20.3 |
| LYS | 860 | A | LYS | 873 | B | 21.4 |
| LYS | 873 | A | LYS | 873 | B | 12.8 |
| LYS | 890 | B | LYS | 901 | B | 16.3 |
| LYS | 890 | A | LYS | 901 | A | 16.4 |
| LYS | 890 | A | LYS | 901 | B | 18.3 |
| LYS | 901 | A | LYS | 901 | B | 9.3 |
| LYS | 901 | A | LYS | 911 | B | 17.9 |
| LYS | 901 | B | LYS | 911 | B | 18.3 |
| LYS | 901 | B | LYS | 920 | B | 20.7 |
| LYS | 901 | A | LYS | 911 | A | 21.8 |
| LYS | 901 | A | LYS | 920 | B | 23.9 |
| LYS | 907 | A | LYS | 911 | B | 7.7 |
| LYS | 907 | A | LYS | 907 | B | 10.6 |
| LYS | 907 | B | LYS | 920 | B | 11.5 |
| LYS | 907 | A | LYS | 911 | A | 12.1 |
| LYS | 907 | B | LYS | 911 | B | 14.4 |
| LYS | 907 | A | LYS | 920 | B | 18.9 |
| SER | 909 | A | LYS | 911 | A | 7.1 |
| SER | 909 | B | LYS | 911 | B | 7.4 |
| SER | 909 | A | LYS | 911 | B | 9.4 |
